# Systematic transcriptomics reveals a biphasic mode of sarcomere morphogenesis in flight muscles regulated by Spalt

**DOI:** 10.1101/229534

**Authors:** Maria L. Spletter, Christiane Barz, Assa Yeroslaviz, Xu Zhang, Sandra B. Lemke, Erich Brunner, Giovanni Cardone, Konrad Basler, Bianca H. Habermann, Frank Schnorrer

## Abstract

Muscles organise pseudo-crystalline arrays of actin, myosin and titin filaments to build force-producing sarcomeres. To study how sarcomeres are built, we performed transcriptome sequencing of developing *Drosophila* flight muscles and identified 40 distinct expression profile clusters. Strikingly, two clusters are strongly enriched for sarcomeric components. Temporal gene expression together with detailed morphological analysis enabled us to define two distinct phases of sarcomere development, which both require the transcriptional regulator Spalt major. During the sarcomere formation phase, 1.8 μm long immature sarcomeres assemble myofibrils that spontaneously contract. During the sarcomere maturation phase, these sarcomeres grow to their final 3.2 μm length and 1.5 μm diameter and acquire stretch-sensitivity. Interestingly, the final number of myofibrils per flight muscle fiber is determined at the onset of the first phase. Together, this defines a biphasic mode of sarcomere and myofibril morphogenesis – a new concept that may also apply to vertebrate muscle or heart development.

## Introduction

Sarcomeres are the stereotyped force producing mini-machines present in all striated muscles of bilaterians. They are built of three filament types arrayed in a pseudocrystalline order: actin filaments are cross-linked with their plus ends at the sarcomeric Z-disc and face with their minus ends towards the sarcomere center. In the center, symmetric bipolar muscle myosin filaments, anchored at the M-line, can interact with the actin filaments. Myosin movement towards the actin plus ends thus produces force during sarcomere shortening. Both filament types are permanently linked by a third filament type, the connecting filaments, formed of titin molecules (Gautel and Djinovic-Carugo, 2016; Lange et al., 2006). A remarkable feature of sarcomeres is their stereotyped size, ranging from 3.0 to 3.4 μm in relaxed human skeletal muscle fibers (Ehler and Gautel, 2008; Llewellyn et al., 2008; Regev et al., 2011). Even more remarkable, the length of each bipolar myosin filament is 1.6 μm in all mature sarcomeres of vertebrate muscles, requiring about 300 myosin hexamers to assemble per filament (Gokhin and Fowler, 2013; Tskhovrebova and Trinick, 2003).

Human muscle fibers can be several centimetres in length and both ends of each fiber need to be stably connected to tendons to achieve body movements. As sarcomeres are only a few micrometres in length, many hundreds need to assemble into long linear myofibrils that span from one muscle end to the other and thus enable force transmission from the sarcomeric series to the skeleton (Lemke and Schnorrer, 2017a). Thus far, we have a very limited understanding of how sarcomeres initially assemble into long immature myofibrils during muscle development to exactly match the length of the mature muscle fiber (Sparrow and Schöck, 2009). In particular, we would like to understand how such sarcomeres mature to the very precise stereotyped machines present in mature muscle fibers.

Across evolution, both the pseudo-crystalline regularity of sarcomeres as well as their molecular components are well conserved (Ehler and Gautel, 2008; Vigoreaux, 2006). Thus, *Drosophila* is a valid model to investigate the biogenesis of sarcomeres as well as their maturation. In particular, the large indirect flight muscles (IFMs) that span the entire fly thorax are an ideal model system to investigate mechanisms of myofibrillogenesis. They contain thousands of myofibrils consisting of 3.2 μm long sarcomeres (Schönbauer et al., 2011; Spletter et al., 2015).

Like all *Drosophila* adult muscles, IFMs are formed during pupal development from a pool of undifferentiated myoblasts called adult muscle precursors (AMPs) (Bate et al., 1991). From 8 h after puparium formation (APF), these AMPs either fuse with themselves (for the dorso-ventral flight muscles, DVMs) or with remodelled larval template muscles (for the dorso-longitudinal flight muscles, DLMs) to form myotubes (Dutta et al., 2004; Fernandes et al., 1991). These myotubes develop dynamic leading edges at both ends and initiate attachment to their respective tendon cells at 12 to 16 h APF (Weitkunat et al., 2014). These attachments mature and mechanical tension is built up in the myotubes, followed by the formation of the first immature periodic myofibrils at 30 h APF when the muscle fibers are about 150 μm in length (Weitkunat et al., 2014). These immature myofibrils contain the earliest sarcomeres, which are about 1.8 μm in length (Weitkunat et al., 2014). During the remaining 3 days of pupal development, the muscle fibers grow to about 1 mm to fill the entire thorax and sarcomere length increases to the final length of about 3.2 μm in adult flies (Orfanos et al., 2015).

After myoblasts have fused to myotubes, the flight muscle specific selector gene *spalt major* (*spalt, salm*) is turned on in the developing flight muscle myotubes. Spalt major is responsible for the correct fate determination and development of the flight muscles, which includes the fibrillar flight muscle morphology and the stretch-activated muscle contraction mode (Schönbauer et al., 2011; Syme and Josephson, 2002). It does so by controlling the expression of more than 700 flight muscle specific genes or gene isoforms during development (Spletter and Schnorrer, 2014; Spletter et al., 2015). However, how the interplay between all these isoforms instructs the formation of highly regular, pseudo-crystalline sarcomeres in the flight muscle is not understood.

Here, we studied the dynamics of flight muscle development in detail. We performed a systematic mRNA-Seq time-course of isolated muscle tissue at 8 time points from the myoblast stage until the mature adult muscle stage. Bioinformatic analysis of expression dynamics identified two gene clusters that are strongly enriched for sarcomeric genes. The temporal dynamics of these clusters enabled us to define two distinct phases of sarcomere morphogenesis: a first sarcomere formation phase in which immature myofibrils with short and thin sarcomeres are built, and a second sarcomere maturation phase during which the immature sarcomeres grow to their final length and diameter and functionally mature. Interestingly, all of the myofibrils assemble simultaneously, with the final number of myofibrils being determined at the beginning of the sarcomere formation phase. During the sarcomere maturation phase myofibril number remains constant, suggesting that every mature sarcomere needs to undergo this biphasic development. Both phases require the activity of the flight muscle selector gene *spalt major*, demonstrating that muscle fiber type-specific transcription is continuously required during all phases of sarcomere formation and maturation. Together, these findings suggest that a precise transcriptional control is required to first assemble and then mature sarcomeres to their pseudo-crystalline regularity.

## Results

### A time-course of indirect flight muscle development

To better understand muscle morphogenesis in general and myofibrillogenesis in particular, we focused on the *Drosophila* indirect flight muscles (IFMs). We hypothesised that major morphological transitions during IFM development may be induced by transcriptional changes, thus we aimed to generate a detailed developmental mRNA-Seq dataset from IFMs. IFMs are built from AMPs that adhere to the hinge region of the wing disc epithelium and are labelled with *Him*-GAL4 driven GFP (Figure 1A) (Soler and Taylor, 2009). At 16 h APF, many of these myoblasts have fused to larval template muscles to build the dorsal-longitudinal flight muscle (DLM) myotubes, which initiate attachment to their tendons. At this stage, the DLM myotubes of fibers 3 and 4 have a length of about 300 μm (Figure 1B). Fusion ceases at about 24 h APF (Figure 1C) and attachment matures until 32 h APF, coinciding with the strong recruitment of βPS-Integrin and the spectraplakin homolog Shortstop (Shot) to the attachment sites. At this stage the myofibers have built up mechanical tension and compacted to a length of about 150 μm, coinciding with the appearance of long Shot-positive tendon extensions that anchor the muscles within the thorax. This important developmental transition is highlighted by the assembly of immature myofibrils visualised by strong F-actin staining throughout the entire muscle fiber (Figure 1D) (Weitkunat et al., 2014).

**Figure 1.**
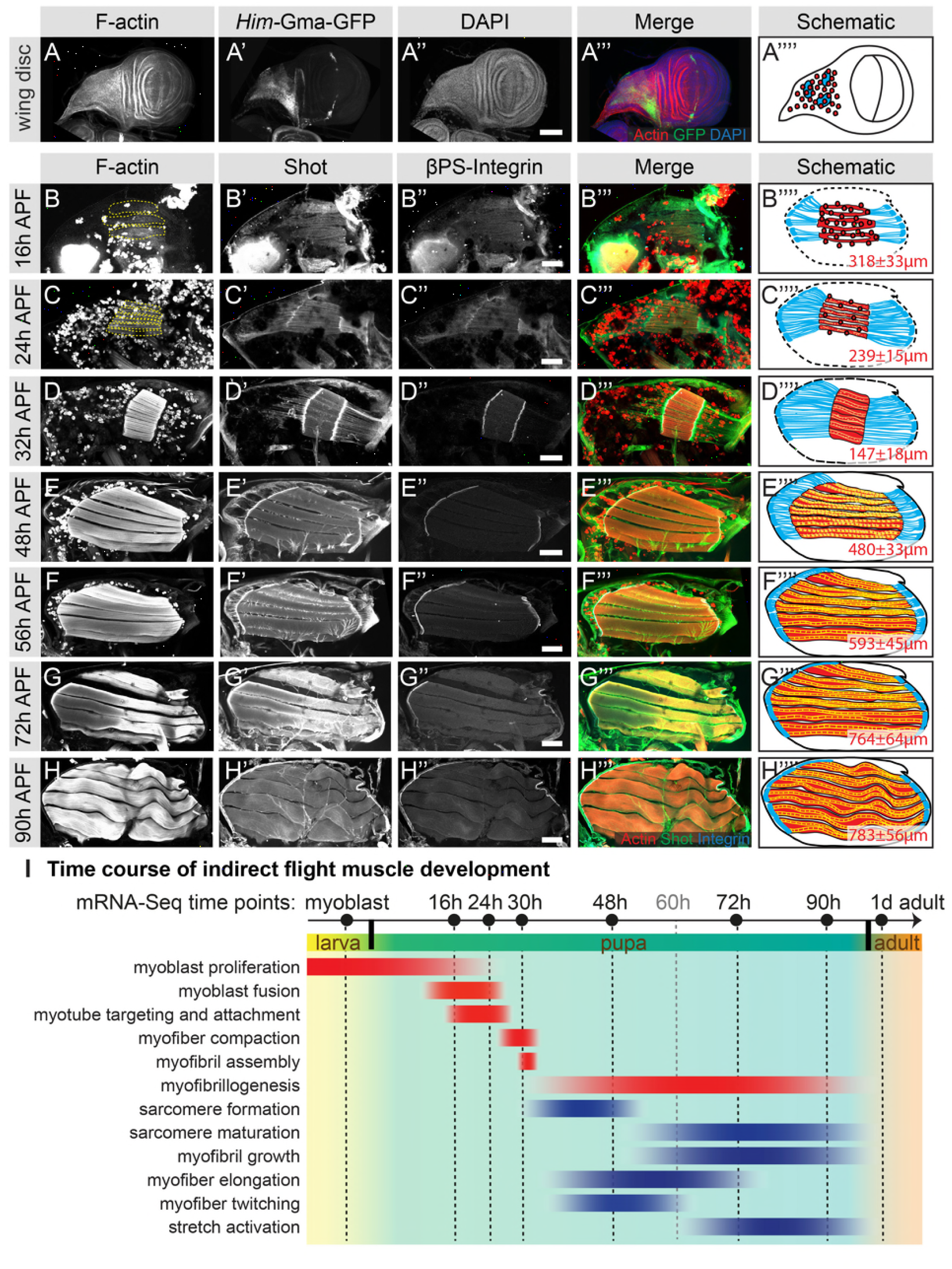
Development of the dorsal longitudinal indirect flight muscles. **(A-H)** Time course of DLM development. (A) Myoblasts adhering to the hinge of the larval wing disc were visualised with *Him*-Gma-GFP (green), F-actin was stained with phalloidin (red) and nuclei with DAPI (blue). (B-H) Time course of DLM and myotendinous junction development at 16h (B), 24h (C), 32h (D), 48h (E), 56h (F), 72h (G) and 90h APF (H). F-actin was stained with phalloidin (red), Shot (green) and βPS-Integrin (blue). For details see text. Scale bar represents 100 μm. (**I**) Temporal summary of known events during myogenesis (red) and events defined in this study (blue). Samples for mRNA-Seq were collected at time points noted in black.

After 32 h APF, the myofibers undergo another developmental transition and begin to grow dramatically. They elongate 3-fold to reach a length of about 480 μm by 48 h APF (Figure 1E) and about 590 μm by 56 h APF (Figure 1F). Concomitantly, the tendon extensions shrink with the myofibers being directly connected to the basal side of the tendon cell epithelium by 72 h APF (Figure 1G). At the end of pupal development (90 h APF), wavy muscle fibers with a length of about 780 μm containing mature myofibrils (Figure 1H) are present within the thorax.

### Expression dynamics during indirect flight muscle development

To quantify transcriptional dynamics across the entire developmental time course, we focused on the major developmental transitions and isolated mRNA from dissociated myoblasts of dissected or mass-isolated third instar wing discs and from hand-dissected IFMs at 16 h, 24 h, 30 h, 48 h, 72 h and 90 h APF pupae, and adult flies 1 day after eclosion (Figure 1I). We performed mRNA-Seq using at least two biological replicates for each time point (see Materials and Methods). To identify genes with similar temporal expression profiles, we used Mfuzz (Kumar and E Futschik, 2007) to cluster standard normalized read counts from all genes expressed above background (12,495 of 13,322 genes). This allowed us to confidently identify 40 distinct genome-wide clusters (Figure 2-S1), each of which contains a unique gene set ranging from 155 to 703 members (Supplementary Table 1). These clusters represent various temporal expression dynamics, with high expression at early (myoblast proliferation and fusion), mid (myotube attachment and myofibril assembly) or late (myofiber growth) myogenesis stages or a combination thereof (Figure 1I, Figure 2-S1). These distinct patterns suggest a precise temporal transcriptional regulation corresponding to observed morphological transition points.

To verify the mRNA-Seq and cluster analysis, we selected a number of ‘indicator’ genes with available antibodies or GFP fusion proteins whose expression correlates with important developmental transitions. Twist (Twi) is a myoblast nuclear marker at larval stages and its expression needs to be down-regulated after myoblast fusion in pupae (Anant et al., 1998). We find *twi* mRNA in Mfuzz cluster 27, with high expression in myoblasts until 16 h APF and a significant down-regulation from 24 h APF, which we were able to verify with antibody stainings (Figure 2A-C). The flight muscle fate selector gene *spalt major* (*salm*) (Schönbauer et al., 2011) and its target, the IFM splicing regulator *arrest* (*aret, bruno*) (Spletter et al., 2015) are members of cluster 26 and 14, respectively. Expression of both clusters is up-regulated after myoblast fusion at 16 h APF, which we were able to verify with antibody stainings (Figure 2D-F, Figure 2-S2A-C). For the initiation of muscle attachment, we selected Kon-tiki (Kon) (Schnorrer et al., 2007; Weitkunat et al., 2014), member of cluster 15, which is transiently up-regulated after myoblast fusion before it is down-regulated again after 30 h APF. Consistently, we found Kon-GFP present at muscle attachment sites at 30 h APF but not at 72 h APF (Figure 2G-I). A similar expression peak shifted to slightly later time points is found in cluster 34, which contains *β-tubulin 60D* (*βTub60D*) (Leiss et al., 1988). Consistently, we find β-Tub60D-GFP (Sarov et al., 2016) expression in IFMs at 30 h and 48 h but not 72 h APF (Figure 2J-L). After attachment is initiated, the attachments need to mature and be maintained. As expected, we found the essential attachment components βPS-Integrin (*mys*) and Talin (*rhea*) in clusters that are up-regulated after myoblast fusion until adulthood (clusters 7 and 25, respectively). This is consistent with continuous high protein expression of βPS-Integrin-GFP and Talin-GFP at muscle attachment sites (Figure 2M-O, Figure 2-S2D-F). Taken together, these semi-quantitative protein localisation data nicely validate the temporal mRNA dynamics found in the mRNA-Seq data, confirming our methodology.

**Figure 2.**
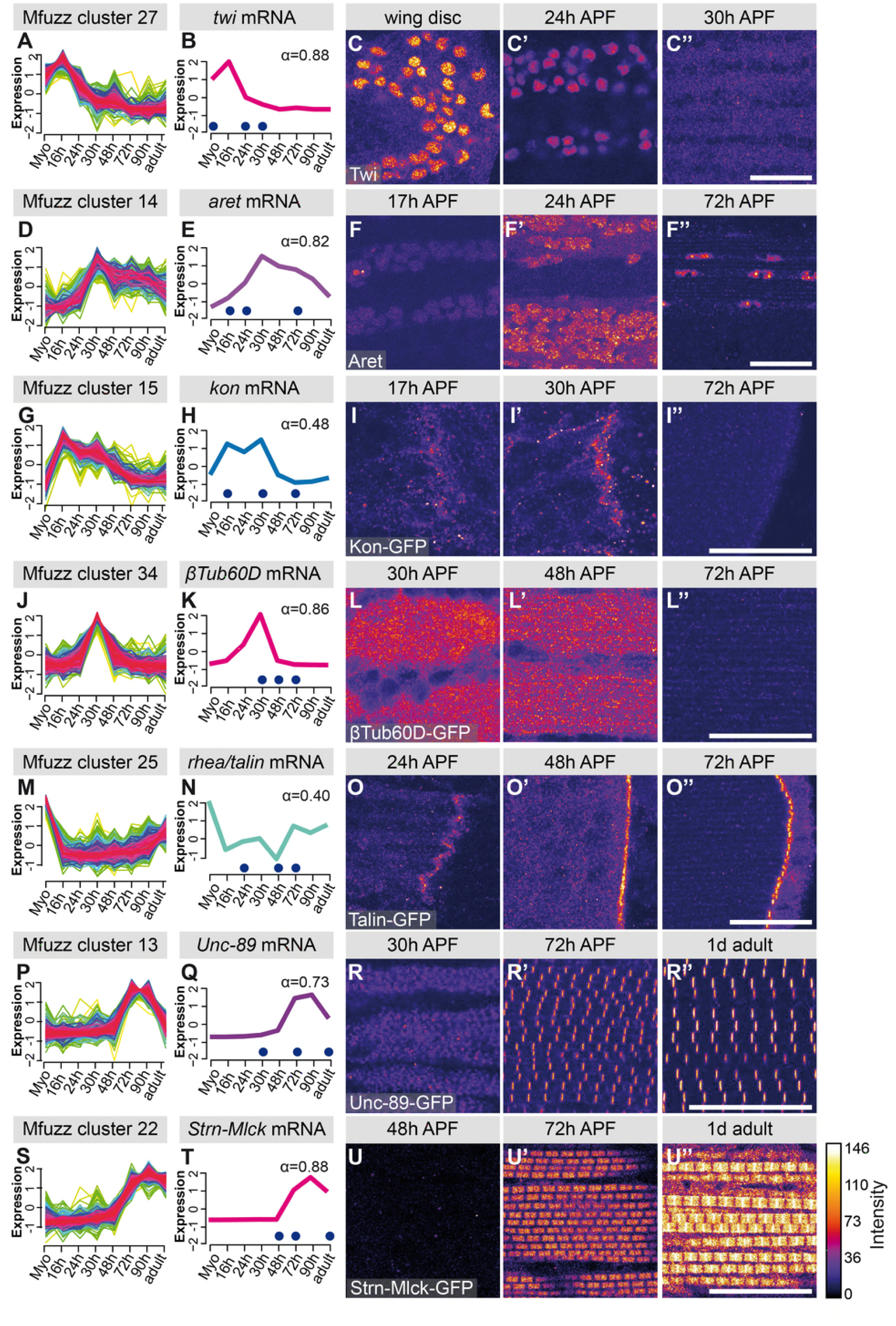
Verification of mRNA-Seq time course by ‘indicator’ gene expression. **(A,D,G,J,M,P,S)** Temporal expression dynamics were evaluated by clustering standard-normal mRNA-seq counts using Mfuzz. Temporal expression profiles are plotted with high membership values in warm colours (red, pink), and lower membership values in cool colours (blue, green). **(B,E,H,K,N,Q,T)** The profile of one ‘indicator’ gene from each cluster is shown and coloured based on the Mfuzz membership value α. **(C,F,I,L,O,R,U)**. Protein expression and localisation dynamics were visualised by antibody staining against Twi (C) and Aret (F) or against GFP for GFP tagged fosmid reporters for Kon (I), β-Tub60D (L), Rhea (Talin) (O), Unc-89 (Obscurin) (R) and Strn-Mlck (U). Images for the same protein were acquired using the same settings, and pseudo-coloured based on intensity. Note the close correlation between mRNA and protein expression dynamics. Time points are indicated by blue dots on the mRNA expression profile. Scale bars represent 20 μm.

### Sarcomeric and mitochondrial gene induction after 30 h APF

Hierarchical clustering of the core expression profiles from the 40 identified Mfuzz clusters defines 8 temporally ordered groups (Figure 3) that show progressive expression dynamics as muscle development proceeds. GO-Elite analysis (Zambon et al., 2012) for gene set enrichment identified gene ontology (GO) terms related to cell proliferation and development as enriched in the early clusters (such as the *twi* cluster 27 or the *kon* cluster 15), whereas terms related to actin filament dynamics are more enriched in the middle clusters (such as the βPS-Integrin cluster 7 and the Talin cluster 25), reassuring that the clustering approach is valid (Figure 3, Supplementary Table 2). Strikingly, the only two clusters that display a strong enrichment for genes important for sarcomere organisation are clusters 13 and 22, both of which are late up-regulated clusters (Figure 3). Members of both clusters just become detectable at 30 h APF (Unc-89/Obscurin-GFP, Act88F-GFP, Mhc-GFP) or even later at 48h APF (Strn-Mlck-GFP, Mf-GFP). In all cases, we could confirm the strong up-regulation from 30 h to 72 h APF in the mRNA-Seq data at the protein level using GFP fusion proteins under endogenous control (Figure 2P-U, Figure 2-S2J-P) (Sarov et al., 2016). We additionally verified the late up-regulation of Flightin (Fln), a member of cluster 3, which is detectable at 72 h but not at 30 h APF (Figure 2-S2G-I).

**Figure 3.**
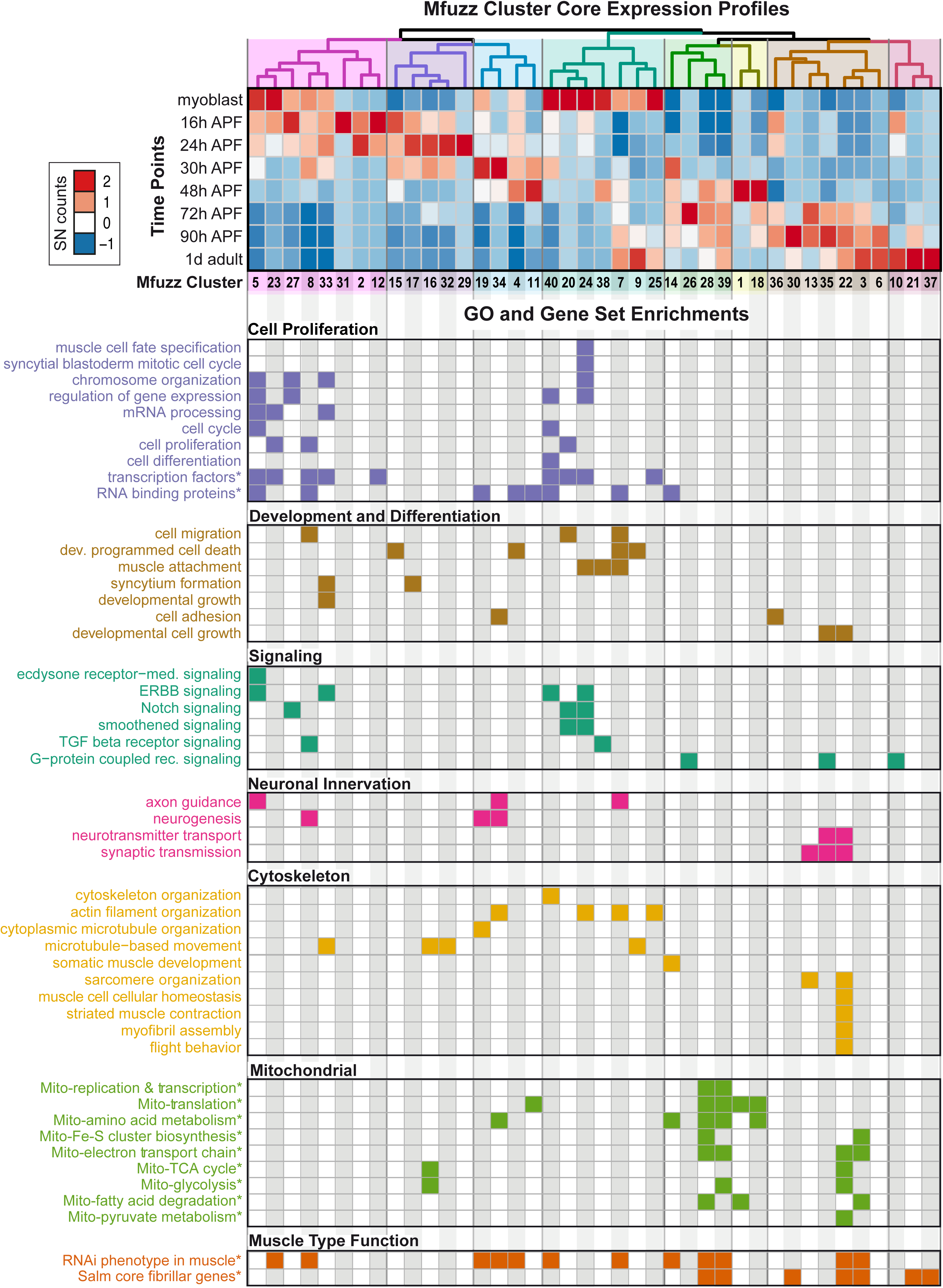
Expression dynamics reveal a temporal ordering of biological processes during muscle morphogenesis. (**Top**) Heat map of Mfuzz cluster core expression profiles. Standard-normal count values for all genes with a membership value α > 0.8 were averaged to generate the core expression profile for each cluster. Mfuzz expression profiles fall into 8 groups (coloured dendrogram leaves) based on hierarchical clustering of their temporal expression dynamics. Time points and Mfuzz clusters are labelled. Colour scale of standard-normal count values ranges from blue (stable/no expression) to red (high expression). (**Bottom**) GO Biological Process and user-defined gene set (marked with *) enrichments calculated with GO-Elite. Note that proliferation, development and differentiation terms are enriched at early time points, while mitochondrial and sarcomere terms are enriched at late time points. A coloured box indicates a significant enrichment of a given term in the specified cluster (see Supplementary Table 2 for details).

At late stages of flight muscle development, mitochondrial density strongly increases (Clark et al., 2006). Using GO-Elite, we found a strong enrichment for mitochondrial related pathways in four late up-regulated clusters, namely 3, 28, 39 as well as the sarcomere cluster 22 (Figure 3). By comparing the clusters to systematic functional data acquired at all stages of *Drosophila* muscle development (Schnorrer et al., 2010), we find enrichments in clusters throughout the time course. Interestingly, genes highly expressed in flight muscle compared to other muscle types, identified as *‘salm*-core genes’ (Spletter et al., 2015), are also enriched in the late clusters, including the mitochondrial enriched clusters 3, 28, 39 and the sarcomere enriched cluster 22 (Figure 3). These data highlight the changes in biological process enrichments that parallel expression dynamics, with a very particular change happening during later stages of muscle development after 30 h APF. This corresponds to a time period after immature myofibrils have been assembled, which thus far remained largely unexplored.

To examine the temporal expression dynamics in more detail, we performed a principle component analysis (PCA) of the mRNA-Seq time-points and Mfuzz clusters and found that the major variance is developmental time, with a notable change after 30 h APF (Figure 4A, Figure 4-S1A). There are a large number of genes being up-regulated as well as down-regulated between 30 h and 48 h and between 48 h and 72 h APF, with major differences between the sets of genes expressed at early (16 h - 30 h APF) versus late (72 h - 90 h APF) stages of development (Figure 4B, Figure 4-S1B). Thus, we focused our attention on the developmental transition between 30 h and 72 h APF, which correlates with major growth of the flight muscle fibers (Figure 1).

**Figure 4.**
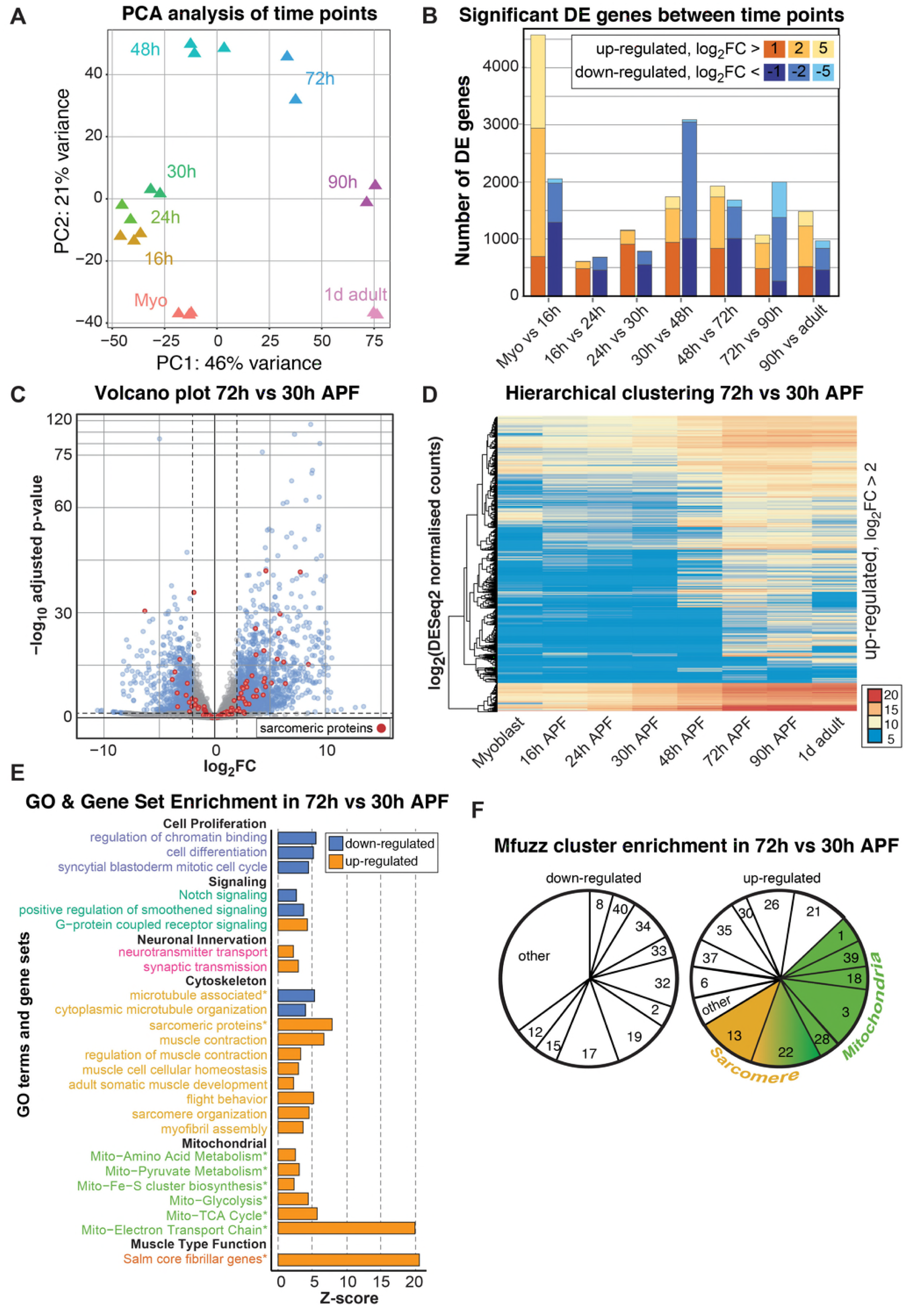
A major induction of sarcomeric gene expression after 30 h APF. **(A)** DESeq2 principle components analysis (PCA) of all mRNA-Seq libraries. Each triangle represents a different biological replicate coloured by time point. Note that individual replicates for a given time point cluster together. PC1 divides early (≤48 h APF) from late (≥72 h) stages. **(B)** Stacked box plot of the number of significantly differentially expressed genes up-regulated (reds) or down-regulated (blues) between sequential time points with a p-value <0.05 and a log_2_FC of >1 (dark), >2 (medium) or >5 (light). The large differences between myoblast to 16 h APF reflect muscle specification. A second large shift in expression is evident between 30 h and 72 h APF. **(C)** Volcano plot illustrating the strong up-regulation of sarcomeric proteins (red) from 30 h to 72 h APF. Significantly up- or down-regulated genes are in blue (p-value <0.05 and abs(log_2_FC) >2). **(D)** Hierarchical clustering of log_2_ transformed DESeq2 normalized counts for all genes that are significantly up-regulated between 30 h and 72 h APF. Note that they are either strongly induced at 48 h or 72 h APF (from yellow to red), or only turned on at 48 h or 72 h APF (blue to yellow/red), suggesting a major transition in gene expression after 30 h APF. Colour scale of log_2_ count values ranges from blue (not expressed) to red (highly expressed). **(E)** GO-Elite enrichments in up- (red) and down- (blue) regulated genes from 30 h to 72 h APF. Note the strong enrichment of mitochondrial and sarcomere terms in the up- regulated genes. **(F)** Pie charts showing the proportion of genes belonging to an enriched Mfuzz cluster in the sets of genes either up- or down-regulated from 30 h - 72 h APF. Note that a large proportion of genes up-regulated 30 h - 72 h belong to cluster 22, as well as Mfuzz clusters enriched for sarcomere (yellow) or mitochondrial (green) terms.

A large number of genes are significantly up- or down-regulated from 30 h to 72 h APF, as visualized on a volcano plot displaying log_2_ fold changes (FC) (Figure 4C), suggesting a major switch in gene expression. In particular, many sarcomeric genes are strongly up-regulated. To identify fine details in expression dynamics, we took all genes significantly up-regulated between 30 h and 72 h APF and performed hierarchical clustering of their DESeq2 normalized counts values (Figure 4D). We noted that many genes are turned on from 30 h to 72 h APF, whereas others are already expressed at 30 h and strongly increase their expression until 72 h (Figure 4D), suggesting two distinct transcriptional phases, before and after 30 h APF. Consistently, we found GO-terms of cell proliferation, cell cycle and Notch signalling down-regulated, whereas actin cytoskeleton, sarcomere, muscle function and mitochondrial related gene sets are strongly up-regulated from 30 h to 72 h APF (Figure 4E). Finally, the genes up-regulated from 30 h to 72 h APF are enriched for sarcomeric proteins and the ‘*salm* core genes’ (Figure 4-S1C, D), and the Mfuzz gene clusters with the most up-regulated members are the mitochondrial and both sarcomeric gene containing clusters 13 and 22 (Figure 4F). Together, these data suggest that in particular the sarcomeric and the mitochondrial genes are strongly induced after 30 h APF.

### A biphasic mode of sarcomere morphogenesis

This interesting strong up-regulation of sarcomeric gene expression after immature myofibrils have been assembled (Figure 1) (Weitkunat et al., 2014) caught our interest and prompted us to more closely investigate the later stages of myofibrillogenesis during which myofibers grow dramatically (Figure 1). We stained the myofibers with phalloidin to reveal myofibril morphology and with the titin isoform Kettin (an isoform of the *sallimus* gene) to label the developing Z-disc and systematically quantified sarcomere length and myofibril width (see Materials and Methods) (Figure 5A, Supplementary Table 3). By measuring the total muscle fiber length, we calculated the total number of sarcomeres per myofibril at a given stage. We found that the sarcomere length and width remain relatively constant at about 2.0 μm and 0.5 μm, respectively until 48 h APF (Figure 5B, C, Supplementary Table 3). However, the number of sarcomeres per myofibril dramatically increases from about 100 at 34 h to about 230 at 48 h APF. After 48 h only a few more sarcomeres are added, resulting in about 270 sarcomeres per myofibril at 60 h APF. This number remains constant until the fly ecloses (Figure 5D, Supplementary Table 3). Moreover, by analysing fiber cross-sections we found that the growth of the individual myofibril diameter correlates with growth of the entire muscle fiber. Both fiber diameter and myofibril diameter remain constant from 30 h to 48 h APF. After 48 h the myofibril diameter grows nearly 3-fold from 0.46 μm to 1.43 μm in adult flies (Figure 5E-G), while fiber cross-sectional area grows nearly 4-fold from 1,759 μm^2^ to 6,970 μm^2^. Strikingly, during the entire time period from 30 h APF to adults, the total number of myofibrils per muscle fiber remains constant (about 2,000 per muscle fiber). Taken together, these quantitative data lead us to propose a biphasic model of sarcomere morphogenesis: 1) During the sarcomere formation phase, which lasts from about 30 h until shortly after 48 h APF, short and thin immature sarcomeres are assembled into immature myofibrils. 2) During the sarcomere maturation phase, starting after 48 h APF, the existing short sarcomeres grow in length and thickness to reach the mature pseudo-crystalline pattern. No new myofibrils are built during the sarcomere maturation phase (Figure 1I).

**Figure 5.**
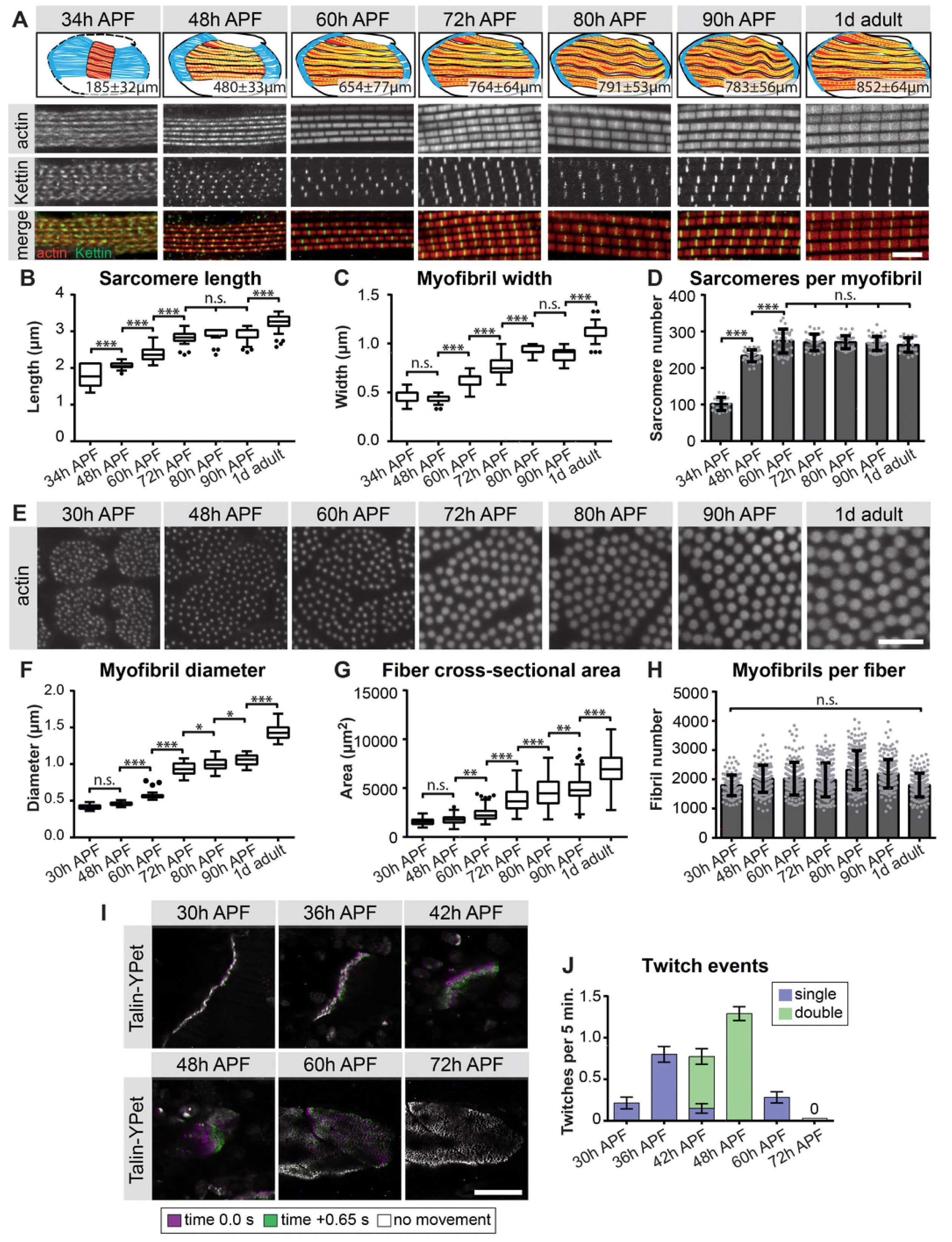
A biphasic mode of sarcomere morphogenesis in flight muscles. **(A)** Scheme of hemi-thoraces at 34 h, 48 h, 60 h, 72 h, 80 h, 90 h APF and 1 day adults (muscle in red, tendon in blue, sarcomeres in yellow) with indicated muscle fiber length. Myofibrils and sarcomeres at these time points were stained for phalloidin (F-actin, red) and Kettin (Z-disc, green). Scale bar represents 5 μm. **(B,C)** Tukey box and whisker plot of sarcomere length and myofibril width. Box extends from 25% to 75%, line marks median, whiskers extend to 25/75% plus 1.5 times the interquartile range. (**D**) Histogram of sarcomere number per myofibril. Error bars represent SD. Note the biphasic mode: sarcomere formation until ~48 h and sarcomere maturation after ~48 h APF. (**E**) Crosssections of the DLMs at 30 h, 48 h, 60 h, 72 h, 80 h, 90 h APF and 1 day adult. Scale bar represents 5 μm. (**F,G**) Tukey box and whisker plot of myofibril diameter and myofiber cross-sectional area. Note the lack of growth in diameter or area from 30 h to 48 h. (**H**) Histogram of the number of myofibrils per myofiber. Error bars represent SD. Note that all myofibrils are already present at 30 h APF and not added later. Tukey’s multiple comparison p-value <.05*, .01**, .001***, n.s. = not significant. N>10 for each individual time point. (**I**) Stills of live movies of DLMs at 30 h, 36 h, 42 h, 48 h, 60 h and 72 h APF. Scale bar represents 50 μm. For live movies see Movie 1. Stills are a time 0.0 s image (magenta) overlaid with a time +0.65 s image (green), where a perfect overlap (white) shows no movement. (**J**) Quantification of spontaneous contraction events per fiber per 5 minutes, with single twitches in blue and double twitches in green. Fibers are first contractile at 30 h APF, reach peak contractility at 48 h and stop all spontaneous contraction shortly after 60 h APF.

We gained additional evidence to support this biphasic assembly model on both the molecular and functional levels. First, the two phases of sarcomere assembly complement the switch in gene expression we observe from 30 h to 72 h APF. Using members of the late induced Mfuzz clusters that contain sarcomeric components, we found that indeed a subset of sarcomeric proteins, such as Unc-89/Obscurin and Mhc, are already detectable in a periodic pattern on immature myofibrils at 30 h. By contrast, other important components, including Fln, Myofilin (Mf) and Strn-Mlck, are only incorporated into myofibrils from 48 h APF, showing high levels by 72 h APF (Figure 5-S1). To functionally test the muscle fibers, we used Talin-YPet as a muscle attachment marker and quantified the number of spontaneous muscle contractions in intact pupae (see Materials and Methods). Interestingly, we found that immature myofibrils already start to spontaneously contract at 30 h APF. These spontaneous contractions increase in strength and frequency until 48 h APF, but then cease, producing no detectable spontaneous contractions at 72 h APF (Figure 5 I,J, Movie 1). This demonstrates that during the sarcomere formation phase, immature contractile myofibrils are generated, which then likely acquire stretch-sensitivity as the immature myofibrils grow and mature during the sarcomere maturation phase, and thus cease contracting.

### Salm induces sarcomeric gene expression during sarcomere maturation

How does biphasic transcription of the various sarcomeric components instruct the biphasic mode of myofibrillogenesis? To address this important question, we performed mRNA-Seq and compared wild type to *spalt-major* knock-down (*salmIR*) flight muscles (Supplementary Table 4). We see down-regulation of mRNAs coding for sarcomeric components at 24 h and 30 h APF, and in particular at 72 h APF during the phase of sarcomere maturation (Figure 6A, Figure 6-S1A-C). The genes down-regulated in *salmIR* IFM are enriched for GO terms associated with sarcomere assembly and flight behaviour and mitochondrial genes, as well as the mitochondrial Mfuzz clusters 3, 28, 39 and the sarcomeric Mfuzz cluster 22 (Figure 6-S1D-F). Indeed, members of cluster 22, which is strongly enriched for sarcomeric and mitochondrial genes, are less strongly induced from 30 h to 72 h APF in *salmIR* muscle compared to wild type (Figure 6B, C), suggesting that *salm* is indeed required for the strong induction of sarcomeric protein expression after 30 h APF.

**Figure 6.**
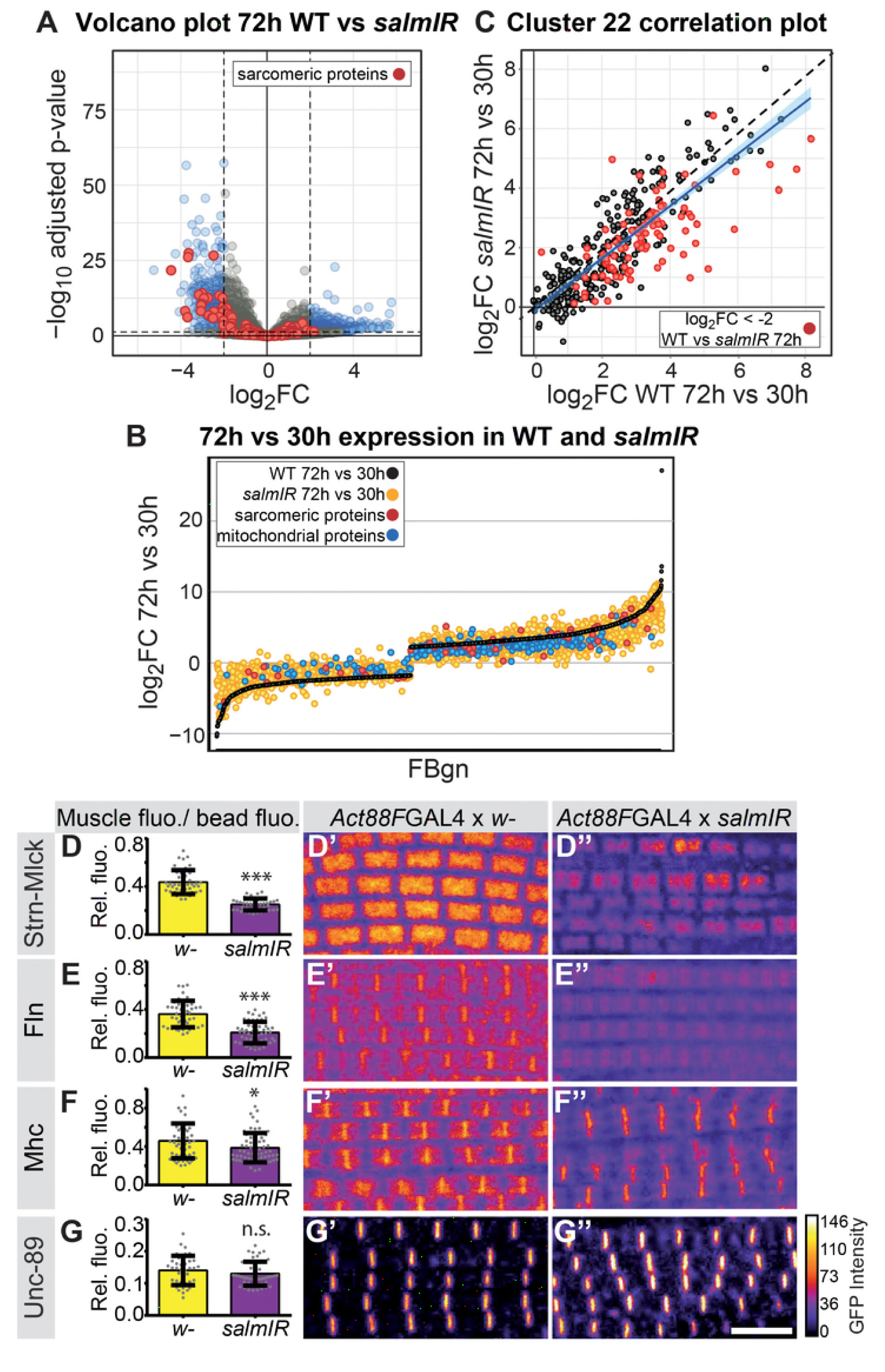
*salm* regulates the transition in gene expression after 30 h APF. **(A)** Volcano plot of mRNA-Seq comparison of wild-type (WT) versus *salmIR* IFMs at 72 h APF. Note the significant down regulation of genes in *salmIR,* especially sarcomeric protein coding genes (red). Significantly differentially expressed (DE) genes (abs(log_2_ FC)>2, p< 0.05) are in blue. (**B**) WT mRNA-Seq fold change values of all genes significantly DE from 30 h to 72 h APF are ordered from lowest to highest (black). The corresponding *salmIR* fold change is shown in yellow. Both sarcomeric (red) and mitochondrial protein coding genes (blue) are less strongly up- or down-regulated in *salmIR* across the 30 h to 72 h transition. (**C**) Correlation plot of genes in Mfuzz cluster 22 comparing the induction of expression from 30 h to 72 h in WT (x-axis) to *salmIR* (y-axis). Note the general spread, indicating a lack of tight regulation in *salmIR* and the large proportion of sarcomeric genes (red) that show reduced induction. A fitted linear regression (blue) reveals a significant decrease in gene expression in *salmIR.* (**D-G**) *salm* is required for the induction of some but not all sarcomeric proteins. *Act88F*≫*salmIR* in the background of GFP-tagged Strn-Mlck-IsoR (D), Fln (E), Mhc (F) and Unc-89 (G). Quantitative changes in live GFP fluorescence at 90 h APF were measured by quantitative confocal microscopy relative to standard fluorescent beads, revealing significant decreases in induction for Strn-Mlck, Fln and Mhc between wild type control (shown in yellow, *Act88F*-GAL4 crossed to *w^1118^)* and *Act88F*≫*salmIR* (shown in purple). Scale bar represents 5 μm. Error bars represent SEM, Student’s t-test p-value <0.05*, 0.001***, n.s. = not significant. N>10 for each individual sample. (D’-G’’) Intensity-coded GFP fluorescence at 90 h APF in confocal images for fixed myofibrils.

Salm is expressed shortly after myoblast fusion and constitutive knock-down of *salm* with *Mef2*-GAL4 results in a major shift of muscle fiber fate (Schönbauer et al., 2011), which may indirectly influence transcription after 30 h APF. Hence, we aimed to reduce Salm levels only later in development, to directly address its role in the second sarcomere maturation phase. To this end, we knocked-down *salm* with the flight muscle specific driver Act88F-GAL4, which is expressed from about 18 h APF and requires *salm* activity for its expression (Bryantsev et al., 2012; Spletter et al., 2015). This strategy enabled us to reduce Salm protein levels at 24 h APF resulting in undetectable Salm levels at 72 h APF (Figure 6-S2). To test if Salm instructs the transcriptional boost of sarcomeric components after 30 h APF, we performed quantitative imaging using unfixed living flight muscles expressing GFP fusion proteins under endogenous control. We used green fluorescent beads to normalise the GFP intensity between different samples. While overall sarcomere morphology is not strongly affected in *Act88F>>salmIR* muscles, we found that the levels of Strn-Mlck, Fln and Mhc are strongly reduced at 72 h as compared to wild-type controls (Figure 6D-G). This suggests that *salm* is indeed required for the expression boost of a number of sarcomeric proteins during the sarcomere maturation phase after 48 h APF.

To investigate the consequences of late *salm* knock-down, we quantified the myofibril and sarcomere morphology throughout the second phase of sarcomere maturation. The myofibrils display a fibrillar morphology, confirming that the early function of Salm to determine IFM fate was unaffected by our late knock-down. At 72 h APF and more prominently at 90 h APF and in adults, *Act88F*≫*salmIR* myofibrils showed actin accumulations at broadened Z-discs (Figure 7A-H), which are often a landmark of nemaline myopathies (Sevdali et al., 2013; Wallgren-Pettersson et al., 2011). The myofibril width was not significantly different in these myofibrils (Figure 7I). However, the sarcomeres displayed a strong defect in sarcomere length growth after 48 h APF in *Act88F*≫*salmIR* muscles (Figure 7J, Supplemental Table 3), with sarcomeres only obtaining a length of 2.84 μm in adult flies, demonstrating that Salm activity is required for normal sarcomere maturation and growth.

**Figure 7.**
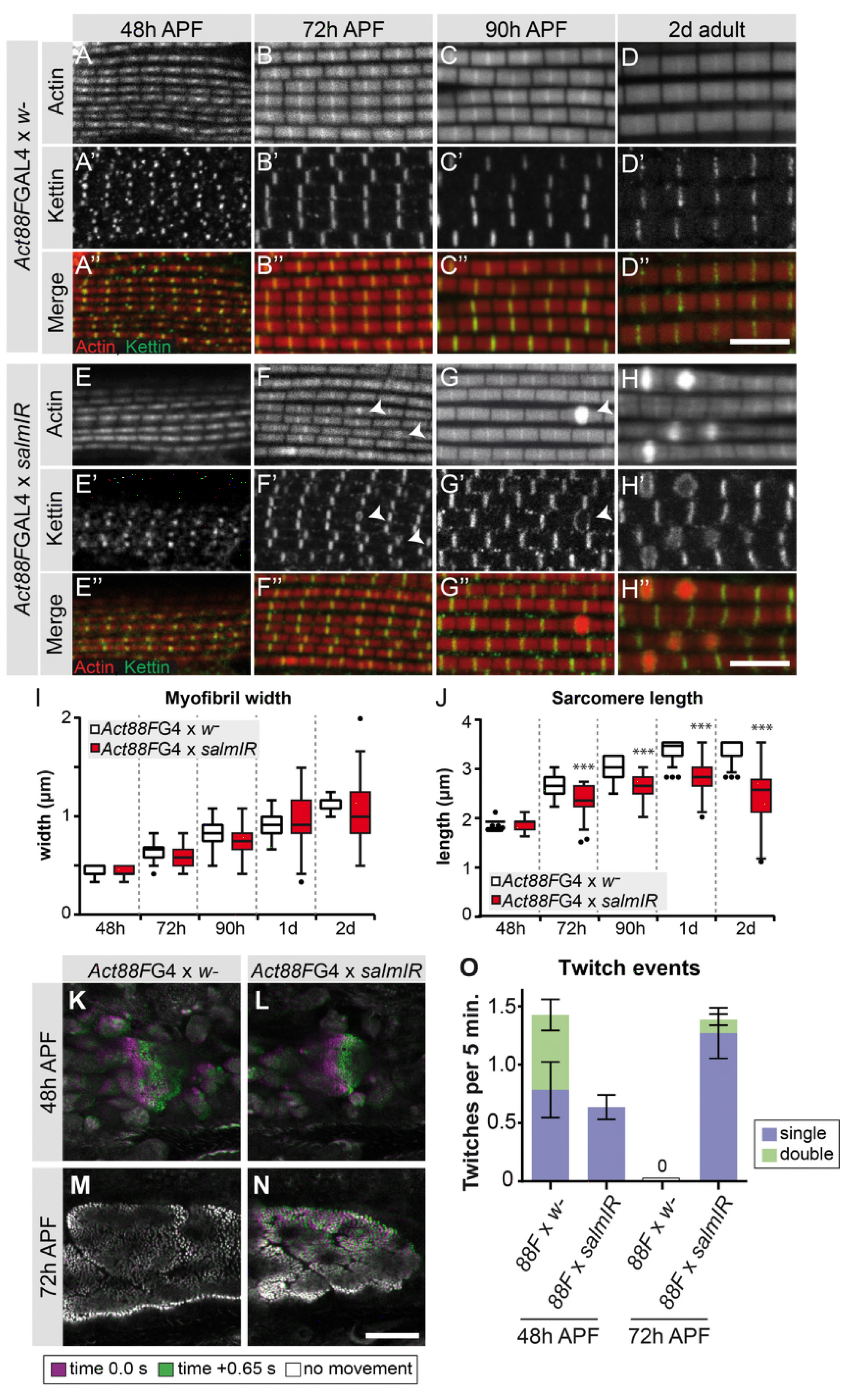
*salm* is required for normal sarcomere maturation and function. **(A-H)** Myofibrils of Act88F-GAL4 / + (A-D) or *Act88F*≫*salmIR* (E-H). Note that *salmIR* DLM remains fibrillar and appears normal at 48 h APF (E), but at 72 h APF (F) 90h APF (G) and 2 day adult (H) Z-discs widen and show actin accumulations (arrowheads). (**I,J**) Tukey box and whisker plot of myofibril width (I) and sarcomere length (J) in Act88F-GAL4 / + and *Act88F*≫*salmIR* (red). Tukey’s multiple comparison p-value <.001***. N>10 for each individual time point. Scale bars represent 5 μm. (**K-O**) Stills from live movies of developing DLMs at 48 h and 72 h APF in Act88F-GAL4 / + (K, M) and *Act88F*≫*salmIR* (L, N). Scale bar represents 50 μm. Coloured as in Figure 5. (O) Quantification of spontaneous contraction events per fiber per 5 minutes, with single twitches in blue and double twitches in green. Error bars represent SEM. *salmIR* fibers continue spontaneously contracting at 72 h APF.

### Salm function contributes to gain of stretch-activation during sarcomere maturation

Given the defects in sarcomere length and sarcomere gene expression in *Act88F*≫*salmIR* muscles, we explored the function of these abnormal muscle fibers. As expected, *Act88F*≫*salmIR* flies are flightless (Figure 7S1A) and we observed rupturing of the adult muscle fibers within 1d after eclosion (Figure 7S1B-G), demonstrating the importance of proper sarcomere maturation to prevent muscle atrophy. Based on our finding that spontaneous flight muscle contractions stop by 72 h APF, we hypothesized that if Salm truly regulates the second phase of sarcomere maturation, we may see spontaneous contraction defects during development. At 48 h APF, *Act88F*≫*salmIR* fibers twitch, but less often than and without the double twitches observed in control fibers (Figure 7K,L,O; Movie 2). Strikingly, at 72 h APF *salmIR* fibers fail to stop contracting and moreover show frequent and uncoordinated spontaneous contractions in which different myofibril bundles of the same fiber twitch at different times (Figure 7M-O, Movie 3), demonstrating that sarcomere maturation is indeed disrupted, with the likely consequence that myofibrils fail to acquire normal stretch-activation sensitivity.

To directly test the function of a sarcomeric component during the sarcomere maturation phase, we investigated the role of the prominently induced Salm target Strn-Mlck, which is largely incorporated during the sarcomere maturation process (Figure 5S1E). In *Strn-Mlck* mutants, sarcomere and myofibril morphology, including myofibril width, is initially normal. However, at 80 h APF the sarcomeres overgrow, consistently reaching lengths of more than 3.5 μm and resulting in slightly longer muscle fibers at 80 h APF (Figure 8). After overgrowing, sarcomeres appear to hyper-contract resulting in short, thick sarcomeres in 1-day-old adults (Figure 8E, J, K, L). Like *Act88F*≫*salmIR* flies, *Strn-Mlck* mutant adults are flightless (Figure 7S1A) and display ruptured fibers during the first days of life (Figure 7S1K-M) (Spletter et al., 2015). Together, these data demonstrate that sarcomere maturation must be precisely controlled at the transcriptional level to enable the precise growth of sarcomeres to their final mature size. This ensures the lifelong function of the contractile apparatus of muscle fibers.

**Figure 8.**
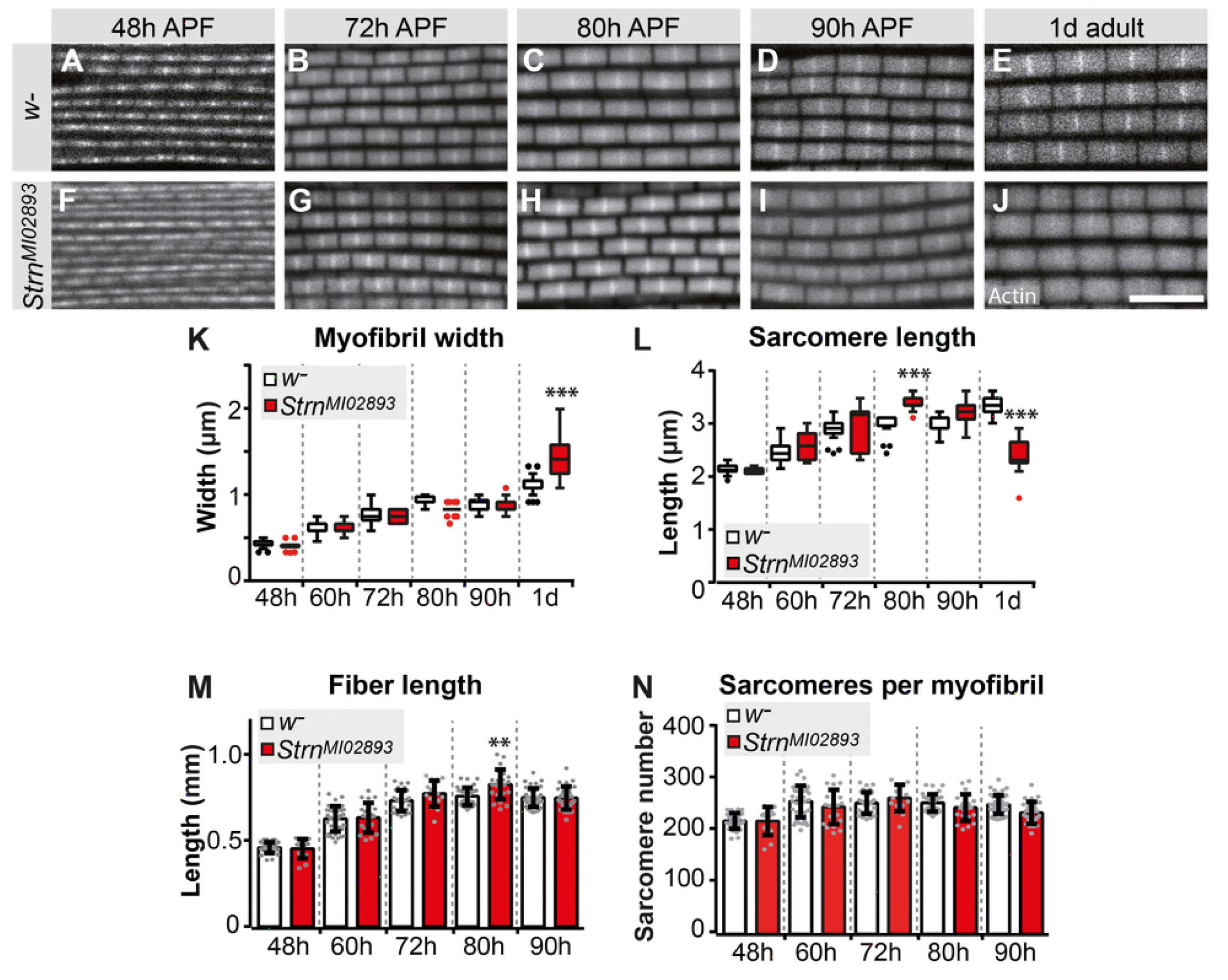
Salm target Strn-Mlck regulates sarcomere length during sarcomere maturation. **(A-J)** Wild type (A-E) and *Strn^M102893^* mutant (F-J) sarcomere development at 48 h, 72 h, 80 h, 90 h APF and 1 day adult. **(K-N)** Tukey box and whisker plot of myofibril width (K) and sarcomere length (L) in wild type and *Strn^M102893^* mutant (red). Tukey’s multiple comparison p-value <.001***. N>10 for each individual time point. Histogram of fiber length (M) and number of sarcomeres per myofibril (N). Error bars represent SEM. Tukey’s multiple comparison p-value <.01**, N>10 for each individual time point. Note that a normal number of sarcomeres are formed in *Strn^M102893^* mutants, but they grow too long at 80 h APF and hyper-contract in 1day adult.

## Discussion

In this study, we generated a systematic developmental transcriptomics resource from *Drosophila* flight muscle. The resource quantifies the transcriptional dynamics across all the major stages of muscle development over five days, starting with stem cell-like myoblasts and attaching myotubes to fully differentiated, stretch-activatable muscle fibers. In this study, we have specifically focused on the transcriptional regulation of sarcomere and myofibril morphogenesis; however, the data we provide cover all other expected dynamics, such as mitochondrial biogenesis, T-tubule morphogenesis, neuromuscular junction formation, tracheal invagination, *etc.* Thus, our data should be a versatile resource for the muscle community.

### A transcriptional switch correlating with two phases of sarcomere morphogenesis

Earlier work has shown that the flight muscle myotubes first attach to tendon cells and then build-up mechanical tension. This tension triggers the simultaneous assembly of immature myofibrils, converting the myotube to an early myofiber (Weitkunat et al., 2014). This suggested a tension-driven self-organisation mechanism of myofibrillogenesis (Lemke and Schnorrer, 2017a). Here we discovered that myofibrillogenesis is not only regulated mechanically, but to a large extent also transcriptionally. At 30 h of pupal development, a large number of genes coding for sarcomeric proteins, including Mhc, Act88F and Unc-89/Obscurin, become up-regulated to enable the first phase of sarcomerogenesis – the assembly of short, immature sarcomeres within thin, immature myofibrils (Figure 9). In this first phase until about 48 h APF, expression of the sarcomeric proteins increases and the muscle fiber grows, accompanied by the elongation of the immature myofibrils through the addition of new sarcomeres. This increases the sarcomere number from about 80 close to the final number of 270 in each myofibril. Thus, we called this first phase the sarcomere formation phase. At its end, the final number of sarcomeres are present within a defined number of myofibrils. These sarcomeres are contractile, but remain short and thin.

**Figure 9.**
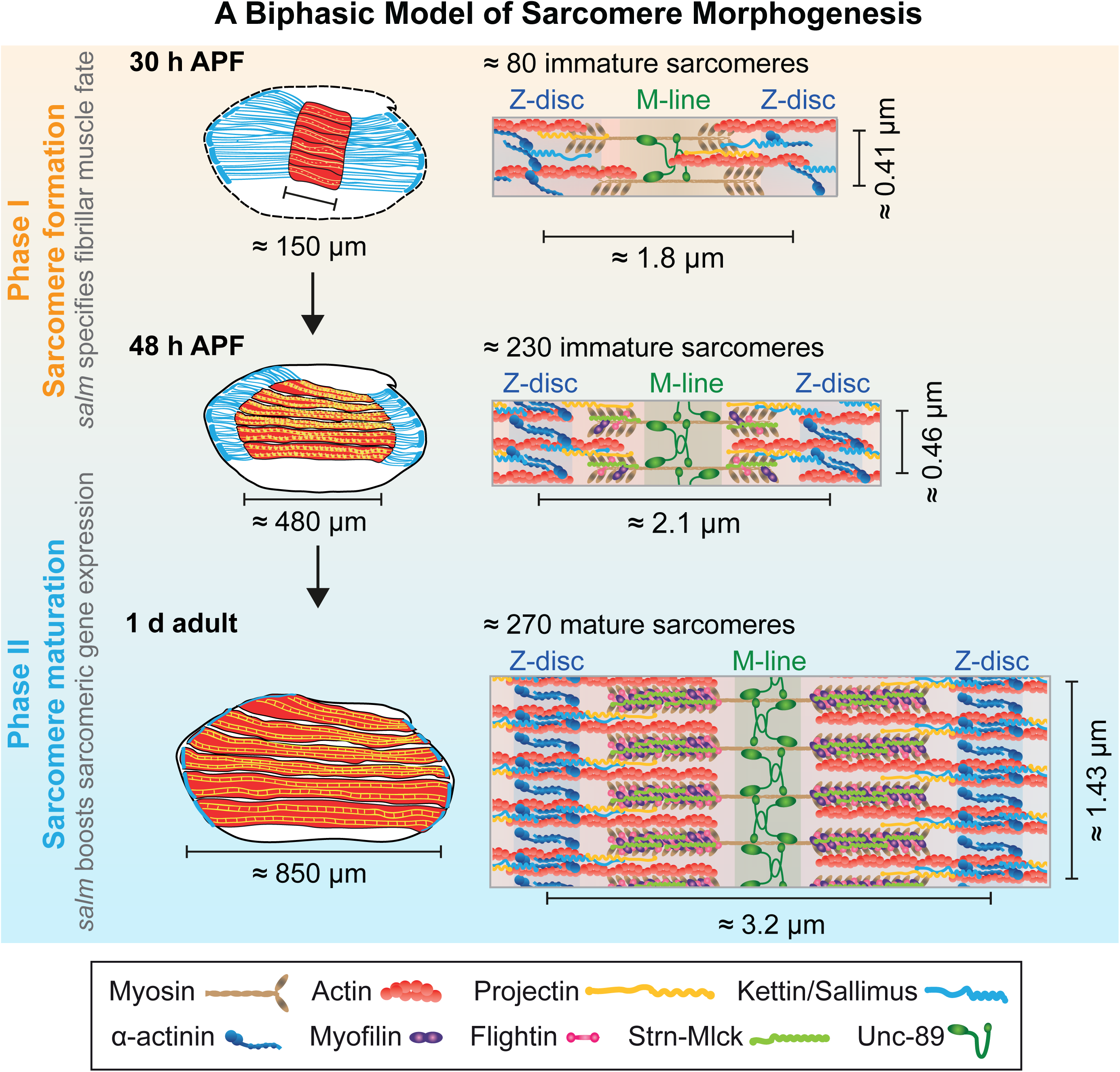
A bi-phasic model of sarcomere morphogenesis. **Phase 1 - Sarcomere formation:** Immature sarcomeres and myofibrils self-assemble around 30 h APF. These immature myofibrils are contractile and increase in length by the addition of new sarcomeres until about 48 h APF. **Phase 2 - Sarcomere maturation:** After 48 h APF, all sarcomeres strongly grow in width and length by the incorporation of new structural proteins. This enables the flight muscle to gain stretch-activation. *salm* is required before the first phase to specify the fibrillar muscle fate and during the second phase to boost the expression of sarcomeric proteins. Muscles are shown in red, tendons in blue. Structural proteins are illustrated as cartoons and are not drawn to scale.

During the second sarcomere maturation phase, the existing immature sarcomeres grow in length and particularly in diameter to reach the pseudo-crystalline regularity of mature myofibrils within about two days of development. This sarcomere maturation phase is initiated by a strong transcriptional burst of the sarcomeric genes. Proteins already present in immature myofibrils like Mhc, Act88F and Unc-89/Obscurin are expressed to even higher levels, and new, often flight-muscle specific proteins like Mf, Fln and the titin-related isoform Strn-Mlck, are expressed to high levels and incorporated into the maturing sarcomeres, facilitating their dramatic growth (Figure 9). Importantly, these matured sarcomeres no longer contract spontaneously, likely because they acquired the stretch-activated mechanism of contraction that is well described for mature *Drosophila* flight muscles (Bullard and Pastore, 2011; Josephson, 2006).

Our biphasic sarcomere morphogenesis model is strongly supported by the observation that the number of sarcomeres per myofibril does not increase after the sarcomere formation phase (ending shortly after 48 h APF). Furthermore, the number of myofibrils remains largely constant during the entire sarcomere morphogenesis period, suggesting that in flight muscles no new myofibrils are added after the initial assembly of immature myofibrils at 32 h APF. Together, this suggests that every sarcomere present in adult flight muscles undergoes this biphasic development.

The model is further supported by previous electron microscopy (EM) studies. We and others found that immature myofibrils have a width of about 0.5 μm (Weitkunat et al., 2014), which corresponds to about 4 thick filaments across each myofibril at the EM level at 42 h APF (at 22°C) (Reedy and Beall, 1993). This ‘core’ myofibril structure built during the sarcomere formation phase is expanded dramatically after 48 h APF, reaching a mature width of 1.5 μm, corresponding to 35 thick filaments across each myofibril at the EM-level (Reedy and Beall, 1993). In total, each adult myofibril contains around 800 thick filaments (Gajewski and Schulz, 2010). The ‘core’ myofibril structure was also revealed by the preferential recruitment of over-expressed actin isoforms (Roper et al., 2005) and more importantly, by selective incorporation of a particular Mhc isoform that is only expressed at mid-stages of flight muscle development (Orfanos and Sparrow, 2013). This Mhc isoform expression switch corresponds to the global switch in sarcomeric gene expression between both sarcomere morphogenesis phases that we defined here. It also fits with recent observations that the formin family member Fhos is important for thin filament recruitment and growth in myofibril diameter after 48 h APF (Shwartz et al., 2016).

### A role of active sarcomere contractions

Mature indirect flight muscles employ a stretch-activated mechanism of muscle contraction, thus Ca^2+^ is not sufficient to trigger muscle contractions without additional mechanical stretch (Bullard and Pastore, 2011; Josephson, 2006). This is different to cross-striated body muscles of flies or mammals that contract synchronously with Ca^2+^ influx. Hence, it is intriguing that immature flight muscle myofibrils do in fact contract spontaneously, with the contraction frequencies and intensities increasing until 48 h APF. It was recently proposed in *Drosophila* cross-striated abdominal muscles and in the developing cross-striated zebrafish muscles that spontaneous contractions are important for the proper formation of the cross-striated pattern (Mazelet et al., 2016; Weitkunat et al., 2017). A similar role for contractions was found in C2C12 cells by stimulating the contractions optogenetically (Asano et al., 2015). This shows that spontaneous contractions are a necessary general feature for the assembly of cross-striated muscle fibers across species.

However, flight muscles are not cross-striated in the classical sense, but have a fibrillar organisation in which each myofibril remains isolated and is not aligned with its neighbouring myofibrils (Figure 5) (Josephson, 2006; Schönbauer et al., 2011). We can only speculate about the mechanism that prevents alignment of the myofibrils in the flight muscles, but it is likely related to their stretch-activated contraction mechanism. This mechanism prevents spontaneous twitching due to increased Ca^2+^ levels, because it additionally requires mechanical activation that can only occur during flight in the adult. Thus, flight muscle sarcomeres not only grow and mature during the second phase of sarcomere maturation, but likely also gain their stretch-activatability.

### Continuous maintenance of muscle type-specific fate

We identified a switch in gene expression between the sarcomere formation and sarcomere maturation phases. Such large scale transcriptome changes have also been observed during mouse (Brinegar et al., 2017), chicken (Zheng et al., 2009) and pig (Zhao et al., 2015) skeletal muscle development or during regeneration after injury in fish (Montfort et al., 2016) and mouse muscles (Warren et al., 2007), indicating that muscle maturation generally correlates with large scale transcriptional changes.

It is well established that general myogenic transcription factors, in particular Mef2, are continuously required in muscles for their normal differentiation (Sandmann et al., 2006; Soler et al., 2012). Mef2 regulates a suit of sarcomeric proteins in flies, fish and mouse muscle important for correct sarcomere assembly and maturation (Hinits and Hughes, 2007; Kelly et al., 2002; Potthoff et al., 2007; Stronach et al., 1999). In *Drosophila,* Mef2 collaborates with tissue-specific factors, such as CF2, to induce and fine-tune expression of structural genes (Gajewski and Schulz, 2010; García-Zaragoza et al., 2008; Tanaka et al., 2008). General transcriptional regulators, such as E2F, further contribute to high levels of muscle gene expression observed during myofibrillogenesis, in part through regulation of Mef2 itself (Zappia and Frolov, 2016). However, it is less clear if muscle type-specific identity genes are continuously required to execute muscle type-specific fate. Spalt major (Salm) is expressed after myoblast fusion in flight muscle myotubes and is required for all flight muscle type-specific gene expression: in its absence the fibrillar flight muscle is converted to tubular cross-striated muscle (Schönbauer et al., 2011; Spletter et al., 2015). Here we demonstrate that Salm is continuously required for correct sarcomere morphogenesis, as late *salm* knock-down leads to defects in sarcomere growth during the sarcomere maturation phase, followed by severe muscle atrophy in adults.

### Two phases of sarcomere morphogenesis – a general mechanism?

Here we defined a biphasic mode of sarcomere morphogenesis in *Drosophila* flight muscles. Is this a general concept for sarcomere morphogenesis? Reviewing the literature, one finds that in other *Drosophila* muscle types which display a tubular crosss-triated myofibril organisation, such as the fly abdominal muscles, the striated sarcomeres also first assemble and then grow in length (Pérez-Moreno et al., 2014; Weitkunat et al., 2017), suggesting a conserved mechanism. In developing zebrafish skeletal muscles, young myofibers present in younger somites show a short sarcomere length of about 1.2 μm, which increases to about 2.3 μm when somites and muscle fibers mature (Sanger et al., 2017; 2009). Interestingly, sarcomere length as well as thick filament length increase simultaneously during fish muscle maturation, indicating that as in flights muscles the length of all sarcomeres in one large muscle fiber is homogenous at a given time (Sanger et al., 2009). Similar results were obtained in mouse cardiomyocytes measuring myosin filament length at young (2 somite) and older (13 somite) stages (Du et al., 2008) and even in human cardiomyocytes, in which myofibrils increase nearly 3fold in width and become notably more organized and contractile from 52 to 127 days of gestation (Racca et al., 2016). These observations strongly suggest that our biphasic sarcomere morphogenesis model is indeed also applicable to vertebrate skeletal and possibly heart muscles. As the progression from phase one to phase two requires a transcriptional switch, it will be a future challenge to identify the possible feedback mechanism that indicates the successful end of phase one or a possible re-entry into the sarcomere formation phase during muscle regeneration or exercise induced muscle fiber growth.

## Materials and Methods

### Fly Strains

Fly stocks were maintained using standard culture conditions. Characterization of normal IFM sarcomere and fiber growth was performed in *w^1118^* grown at 27°C. *salm* RNAi was performed with previously characterized GD3029 (referred to as *salmIR*) and KK181052 (Schönbauer et al., 2011) from VDRC (http://stockcenter.vdrc.at) at 25°C using *Act88F*-GAL4 to induce knock-down after 24 h APF. *Act88F*-GAL4 x *w^1118^* served as control. The *Strn-Mlck-MiMIC* insertion MI02893 into IFM-specific IsoR (Bloomington stock 37038) and TRiP hairpin JF02170 were obtained from Bloomington. The salm-EGFP line was used to sort wing discs (Marty et al., 2014). Tagged genomic fosmid reporter fly lines include strn4 *(Strn-Mlck-GFP*, Isoform R) (Spletter et al., 2015), fTRG500 (*Mhc-GFP,* Isoforms K, L, M), fTRG501 (*Mf-GFP,* Isoforms A, G, N), fTRG587 (*Rhea-GFP,* Isoforms B, E, F, G), fTRG876 (*Fln-GFP*), fTRG932 (*mys-GFP*), fTRG958 (*βTub60D-GFP*), fTRG1046 (*unc-89-GFP),* and fTRG10028 (*Act88F-GFP*) (Sarov et al., 2016).

To label myoblasts, we utilized the enhancer for *Holes-in-muscle* (*Him*), which is expressed in dividing myoblasts and promotes the progenitor fate. *Him-nuc-eGFP* flies were a gift of M. Taylor (Soler and Taylor, 2009). Him-Gal4 flies were created by cloning an EcoRI to SacII fragment of the *Him* enhancer (Liotta et al., 2007) upstream of GAL4 into pStinger. UAS-BBM (UAS-palmCherry) (Förster and Luschnig, 2012) was driven with *Him*-Gal4 to label myoblasts. Him-Gma-GFP flies were created by PCR amplifying Gma-GFP with AscI and PacI overhangs and then cloning downstream of the Him enhancer in pStinger to generate a gypsy insulator-*Him^enh^*-Gma-GFP-SV40-gypsy insulator cassette.

The *rhea-YPet* line used to label muscle ends for live imaging of twitch events was generated by CRISPR-mediated gene editing at the endogenous locus (S.B.L & F.S., details will be published elsewhere). The *kon-GFP* line was generated by inserting GFP into the *kon* locus after its transmembrane domain using the genomic fosmid FlyFos021621, which was integrated using Φ-C31 into VK00033 (I. Ferreira and F.S., details will be published elsewhere).

### Flight tests

Flight tests were performed as previously described (Schnorrer et al., 2010). *Act88F*-GAL4 crosses were kept at 25°C, as higher temperatures negatively impacted flight ability, because of the very high GAL4 expression levels in this strain. Adult males were collected on CO_2_ and recovered at least 24 h at 25 °C before testing. Flies were introduced into the top of a 1 m long cylinder divided into 5 zones. Those that landed in the top two zones were considered ‘normal fliers’, those in the next two zones ‘weak fliers’ and those that fell to the bottom of the cylinder ‘flightless’.

### Immuno-staining

Wing-discs were dissected from 3^rd^ instar wandering larvae in 1x PBS and fixed in 4% PFA in PBS-T. Discs were stained as described below for anti-GFP. Adult and pupal flight muscles were dissected and stained as previously described (Weitkunat and Schnorrer, 2014). Briefly, early pupae (16 h - 60 h APF) were freed from the pupal case, fixed for 20 min. in 4% PFA in relaxing solution and washed in 0.5% PBS-Triton-X100 (PBS-T). 72 h APF and older samples were cut sagittally with a microtome blade. All samples were blocked for at least 1 hour at RT in 5% normal goat serum in PBS-T and stained with primary antibodies overnight at 4°C. Primary antibodies include: guinea pig anti-Shot 1:500 (gift of T. Volk), rat anti-Kettin 1:50 (MAC155/Klg16, Babraham Institute), rabbit anti-GFP 1:1000 (ab290, Abcam), rat anti-Bruno 1:500 (Filardo and Ephrussi, 2003), rabbit anti-Salm 1:50 (Kühnlein et al., 1994), mouse anti-βPS-integrin 1:500 (CF.6G11, DSHB), rabbit anti-Twi 1:1000 (gift of Siegfried Roth) and rabbit anti-Fln 1:50 (Reedy et al., 2000)(gift of Jim Vigoreaux). Samples were washed three times in 0.5% PBS-T and incubated overnight at 4°C with secondary conjugated antibodies (1:500) from Invitrogen (Molecular Probes) including: Alexa488 goat anti-guinea pig IgG, Alexa488 donkey anti-rat IgG, Alexa488 goat anti-mouse IgG, Alexa488 goat antirabbit IgG, rhodamine-phalloidin, Alexa568 goat anti-rabbit IgG and Alexa633 goat antimouse IgG. Samples were washed three times in 0.5% PBS-T and mounted in Vectashield containing DAPI.

### Cryosections

Head, wings and abdomen were removed from one day old *w^1118^* flies and thoraxes were fixed overnight at 4°C in 4% PFA. For 30 - 90 h APF samples, pupae were freed from the pupal case, poked 3-5 times with an insect pin in the abdomen and fixed overnight at 4°C in 4% PFA. Thoraxes or pupae were then sunk in 30% sucrose in 0.5% PBS-T overnight at 4°C on a nutator. Thoraxes or pupae were embedded in Tissue-Tek O.C.T. (Sakura Finetek) in plastic moulds (#4566, Sakura Finetek) and frozen on dry ice. Blocks were sectioned at 30 μm on a cryostat (Microm vacutome). Sections were collected on glass slides coated with 1% gelatin + 0.44 μM chromium potassium sulfate dodecahydrate to facilitate tissue adherence. Slides were post-fixed for 1 min. in 4% PFA in 0.5% PBS-T at RT, washed in 0.5% PBS-T, incubated with rhodamine-phalloidin for 2 hours at RT, washed three times in 0.5% PBS-T and mounted in Fluoroshield with DAPI (#F6057, Sigma).

### Microscopy and image analysis

Images were acquired with a Zeiss LSM 780 confocal microscope equipped with an α Plan-APOCHROMAT 100x oil immersion objective lens (NA 1.46). To compare if indicator protein expression replicates the mRNA-Seq expression dynamics, we imaged three time points from each expression profile with the same confocal settings. Laser gain and pinhole settings were set on the brightest sample and reused on remaining time points in the same imaging session. All samples were additionally stained with the same antibody mix on the same day and if possible in the same tube. Images were processed with Fiji (Schindelin et al., 2012) and Photoshop, and displayed using the ‘Fire’ look-up table.

Fiber length and fiber cross-sectional area were measured with freehand drawing tools in Fiji based on rhodamine-phalloidin staining. Sarcomere length, myofibril width, and myofibril diameter were measured automatically using a custom Fiji plug-in, MyofibrilJ, available from https://imagej.net/MyofibrilJ. All measurements are based on rhodamine-phalloidin staining, except 34 h APF sarcomere lengths, which are based on both rhodamine-phalloidin and Unc-89-GFP staining. ‘Sarcomeres per fibril’ was calculated as average individual fiber length divided by sarcomere length for fiber 3 or 4. ‘Fibrils per fiber’ was calculated as average number of fibrils per unit area multiplied by individual fiber cross sectional area.

For determining myofibril diameter, samples were imaged using a 3x optical zoom (50 nm pixel size). At least 20 cross-section images from different fibers for >10 flies were acquired for each time point. The number of fibrils per section and fibril diameter were determined with the tool ‘analyze myofibrils crosswise’ from MyofibrilJ. In this tool, an initial estimate of the diameter is obtained by finding the first minimum in the radial average profile of the autocorrelation (Goodman, 1968) of the image. This estimate is used to calibrate the optimal crop area around all the cross-sections in the image, their position previously detected by finding the local intensity peaks. All of the detected cross sections are then combined to obtain a noise-free average representation of the fibril section. Finally, the diameter is calculated by examining the radial profile of the average and measuring the full width where the intensity is 26% of the maximum range.

For determining sarcomere length and myofibril width, for each experiment between 10 and 25 images were acquired from more than 10 individual flies. From each image, 9 non-overlapping regions of interest were selected, which were rotated to orient fibrils horizontally, when necessary. The tool ‘analyze myofibrils lengthwise’ from MyofibrilJ reports the sarcomere length (indicated as repeat) and myofibril width (indicated as thickness). Because of the periodic nature of sarcomere organization, their length is estimated by means of Fourier analysis, identifying the position of the peaks on the horizontal axis of the Fourier transformed imaged. Quality of the estimate was evaluated by visual inspection of the Fourier transformed image, overlaid with the peaks detected, as generated by the plug-in. Myofibrils width is estimated from the position of the first minimum in the vertical intensity profile of the autocorrelation of the image.

Live imaging of developmental spontaneous contractions was performed on a Leica SP5 confocal microscope. Prior to imaging, a window was cut in the pupal case, and pupae were mounted in slotted slides as previously described (Lemke and Schnorrer, 2017b; Weitkunat and Schnorrer, 2014). At the specified developmental time point, IFMs were recorded every 0.65 seconds for 5 min. General movement within the thorax was distinguished from IFM-specific contraction, and each sample was scored for the number of single or double contractions observed per 5 minute time window. Data were recorded in Excel and ANOVA was performed in GraphPad Prism to determine significant differences. Movies were assembled in Fiji (Image J), cropped and edited for length to highlight a selected twitch event.

Quantitative imaging of fosmid reporter intensity was performed at 90 h APF in live IFM by normalizing to fluorescent beads (ThermoFisher (Molecular Probes), InSpeck™ Green Kit I-7219). IFMs were dissected from 5 flies, mounted with fluorescent microspheres (0.3% or 1% relative intensity, depending on the reporter intensity) in the supplied mounting medium and immediately imaged (within 20 minutes). Intensity measurements were obtained at 40x for at least 10 flies in regions where both IFM and at least 3 beads were visible. Control *Act88F-GAL4* x *w^1118^* and RNAi *Act88F*-GAL4;; *fosmid-GFP* x *salmIR* (fosmids used include *Strn-Mlck-GFP, Mhc-GFP, Fln-GFP,* and *Unc-89-GFP)* were imaged in the same imaging session. Relative fluorescence fiber to beads was calculated for each image in Fiji by averaging intensity for 3 fiber ROIs and 3 bead ROIs. Data were recorded in Excel and Student’s t-test for significance and plotting were performed in GraphPad Prism.

### mRNA-Seq

We previously published mRNA-Seq analysis of dissected IFMs from *Mef2*-GAL4, UAS-GFP-Gma x *w^1118^* at 30 h APF, 72 h APF and 1d adult, and *Mef2-GAL4, UAS-GFP-Gma* x *salmIR* in 1d adult (Spletter et al., 2015). We expanded this analysis in the present study to include myoblasts from 3^rd^ instar larval wing discs (see below) and dissected IFMs from Mef2-GAL4, UAS-GFP-Gma x *w^1118^* at 16 h, 24 h, 30 h, 48 h, 72 h, 90 h APF and from 1 day adults as well as IFMs from Mef2-GAL4, UAS-GFP-Gma x *salmIR* flies at 24 h, 30 h, 72 h APF and from 1 day adults. IFMs were dissected from groups of 15 flies in 30 min to minimize changes to the transcriptome, spun down in PBS for 5 min at 7500 rpm and immediately frozen in 100 μl TriPure reagent (#11667157001, Roche) on dry ice. RNA was isolated after combining IFMs from 150-200 flies, with biological duplicates or triplicates for each time point.

Poly(A)+ mRNA was purified using Dynabeads (#610.06, Invitrogen) and integrity was verified on a Bioanalyzer. mRNA was then fragmented by heating to 94°C for 210 sec in fragmentation buffer (40 mM TrisOAc, 100 mM KOAc, 30 mM MgOAc_2_). First-strand cDNA synthesis was performed with the Superscript III First-Strand Synthesis System (#18080-051, Invitrogen) using random hexamers. The second strand was synthesized with dUTP and submitted to the Vienna Biocenter Core Facilities (VBCF, http://www.vbcf.ac.at) for stranded library preparation according to standard Illumina protocols and sequenced as SR100 on an Illumina HiSeq2500. Libraries were multiplexed two to four per lane using TrueSeq adaptors.

### Wing disc sorting and myoblast isolation

To perform mRNA-Seq on fusion competent myoblasts that will form the IFMs, we first dissected wing discs from wandering 3^rd^ instar larvae and manually cut the hinge away from the wing pouch. mRNA was isolated in TriPure reagent and sequenced as described above. We estimate this sample (Myo1) is ~50% myoblast, as the myoblasts form a nearly uniform layer over the underlying epithelial monolayer. To obtain a purer myoblast sample, we performed large-scale imaginal disc sorting followed by dissociation. We used particle sorting to isolate imaginal discs from *Him*-GAL4, UAS-BBM (UAS-palmCherry); *salm*-EGFP flies based on the green fluorescent signal. 10-12 mL of larvae in PBS were disrupted using a GentleMACS mixer (Miltenyl Biotec) and discs were collected through a mesh sieve (#0278 in, 25 opening, 710 μm). Fat was removed by centrifugation for 10 min. at 1000 rpm at 4°C, discs were rinsed in PBS and then re-suspended in HBSS. Discs were further purified on a Ficoll gradient (25%:16%). Discs were then sorted on a Large Particle Flow Cytometer (BioSorter (FOCA1000), Union Biometrica, Inc.), obtaining 600-1000 discs per sample. Discs were spun for 5 min at 600 rcf in a Teflon Eppendorf tube and then re-suspended in the dissociation mixture (200 μl of 10x Trypsin, 200 μl HBSS, 50 μl collagenase (10 mg/mL), 50 μ! dispase (10 mg/mL)). The tube was incubated for 10 min. at RT and then transferred to a thermal shaker for 30 min. at 25°C at 650 rpm. Myoblasts were filtered through a 35 μm tube-cap filter and spun at 600 rcf for 5 min. to pellet the cells. Cells were resuspended in HBSS for evaluation or frozen in TriPure reagent for RNA extraction. We obtained samples with ~90% purity based on counting the number of red fluorescent cells / non-fluorescent + green fluorescent cells in 3 slide regions. mRNA was isolated in TriPure reagent and sequenced as described above, generating the Myo2 and Myo3 samples.

### Analysis of RNA-Seq data

FASTA files were de-multiplexed and base called using Illumina software. Reads were trimmed using the FASTX-toolkit. Sequences were mapped using STAR (Dobin et al., 2013) to the *Drosophila* genome (BDGP6.80 from ENSEMBL). Mapped reads were sorted and indexed using SAMtools (Li et al., 2009), and then bam files were converted to bigwig files. Libraries were normalized based on library size and read-counts uploaded to the UCSC Browser for visualization.

Mapped sequences were run through featureCounts (Liao et al., 2014) and differential expression analysis was performed on the raw counts using DESeq2 (Love et al., 2014). We performed all pairwise comparisons across the time-course as well as between wild-type and *salmIR* samples (Supplementary Tables 1 and 4). All original and processed data can be found as supplemental data or in the Gene Expression Omnibus submission (accession number GSE107247). R packages employed in the analysis include ComplexHeatmap (Gu et al., 2016), CorrPlot (https://github.com/taiyun/corrplot), VennDiagram (Chen, 2016), plyr (Wickham, 2011), reshape2 (Wickham, 2007), ggplot2 (Wickham, 2009) and RColorBrewer (Neuwirth, 2015).

Genome-wide soft clustering was performed in R with Mfuzz (Futschik and Carlisle, 2005), using the DESeq2 normalized count values. We filtered the dataset to include all genes expressed at one time point or more, defining expression as >100 counts after normalization. We then set all count values <100 to 0, to remove noise below the expression threshold. DESeq2 normalized data was standardized in Mfuzz to have a mean value of zero and a standard deviation of one, to remove the influence of expression magnitude and focus on the expression dynamics. We tested “k” ranging from 10-256. We then performed consecutive rounds of clustering to obtain 3 independent replicates with similar numbers of iterations, ultimately selecting a final k=40 clusters with iterations equal to 975, 1064 and 1118. We calculated a “stability score” for each cluster by calculating how many genes are found in the same cluster in each run (Supplementary Table 1). Figures are from the 1064 iterations dataset. Mfuzz cluster core expression profiles were calculated as the average standard-normal expression of all genes with a membership value greater than or equal to 0.8, and then core profiles were clustered in R using Euclidean distance and complete linkage.

Enrichment analysis was performed with GO-Elite (Zambon et al., 2012) using available Gene Ontology terms for *Drosophila.* We additionally defined user provided gene lists for transcription factors, RNA binding proteins, microtubule associated proteins, sarcomeric proteins, genes with an RNAi phenotype in muscle (Schnorrer et al., 2010), mitochondrial genes ( ttp://mitoXplorer.biochem.mpg.de) and *salm* core fibrillar genes (Spletter et al., 2015). Full results and gene lists are available in Supplementary Table 2. These user-supplied lists allowed us to define more complete gene sets relevant to a particular process or with a specific localization than available in existing GO terms. Analysis was performed with 5000 iterations to generate reliable significance values.

## Data availability

Processed data from DESeq2, Mfuzz and GO-Elite are available in Supplementary Tables 1, 2, 4. mRNA-Seq data are publicly available from NCBI’s Gene Expression Omnibus (GEO) under accession number GSE107247. Fiji scripts for analysis of sarcomere length, myofibril width and myofibril diameter are available from https://imagej.net/MyofibrilJ.

## Acknowledgements

We thank Irene Ferreira for constructing the tagged *kon-tiki* allele and the Bloomington and VDRC stock centers for fly stocks. We are grateful to Reinhard Fässler and Andreas Ladurner for generous support and to Bettina Stender for excellent technical assistance. We thank Olena Nikonova for initial analysis of the *Act88F*-GAL4 x *salmIR* phenotype and Sandra Esser for testing the *salm* KK181052 hairpin. We acknowledge the VBCF (Vienna, AT) for mRNA-sequencing and the Core Facility Bioimaging at the Max Planck Institute for Biochemistry and the LMU Biomedical Center (Martinsried, DE) for help with confocal image analysis. We thank Florian Marty for help with the BioSorter, and Alexander Stark for help setting-up the mRNA-sequencing. We thank Aynur Kaya-Copur and Wouter Koolhaas for helpful discussions, and Nuno Luis and Vincent Loreau for insightful comments on the manuscript. Our work was supported by the Max Planck Society and the CNRS; postdoctoral Humboldt, EMBO long-term (688-2011), and NIH-NRSA (5F32AR062477) fellowships (M.L.S.), the Frederich-Bauer Stiftung (M.L.S.); a Career Development Award from the Human Frontier Science Program (F.S.), the EMBO Young Investigator Program (F.S.) and the European Research Council under the European Union’s Seventh Framework Programme (FP/2007-2013) / ERC Grant 310939 (F.S), the excellence initiative Aix-Marseille University AMIDEX (F.S.), the ANR-ACHN (F.S.) and the LabEX-INFORM (F.S.).

## Author contribution

M.L.S. performed most of the experiments with important support by C.B.. A.Y., M.L.S. and B.H.H. performed the bioinformatic data analysis. X.Z. contributed to analysis of the IFM developmental time course. S.B.L. performed live-imaging and quantification of fiber contractions. E.B. and K.B. generated flies and developed wing-disc sorting. G.C. developed the MyofibrilJ scripts. F.S. conceived and supervised the project. M.L.S. and F.S. made the figures and wrote the manuscript.

---

**Figure 2 - Supplement 1.**
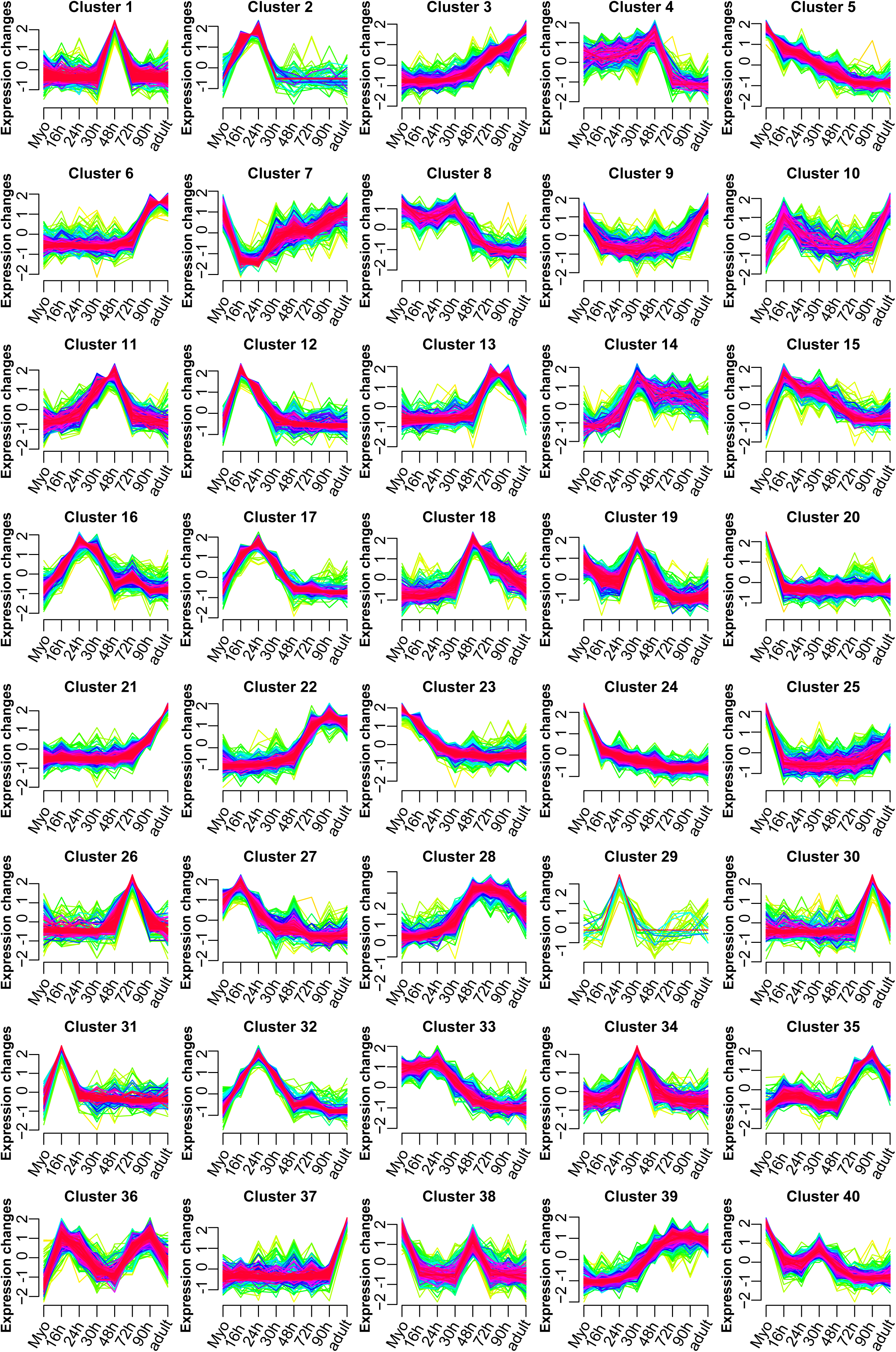
Forty distinct temporal mRNA-Seq expression profiles. Temporal expression profiles were identified by clustering standard-normal mRNA-Seq counts using Mfuzz to group genes with similar temporal expression dynamics. Expression dynamic profiles were labelled 1-40. Each plot shows the profile for each gene in the cluster, with profiles of genes with high membership values in warm colours (red, pink) and lower membership values in cool colours (blue, green).

**Figure 2 - Supplement 2.**
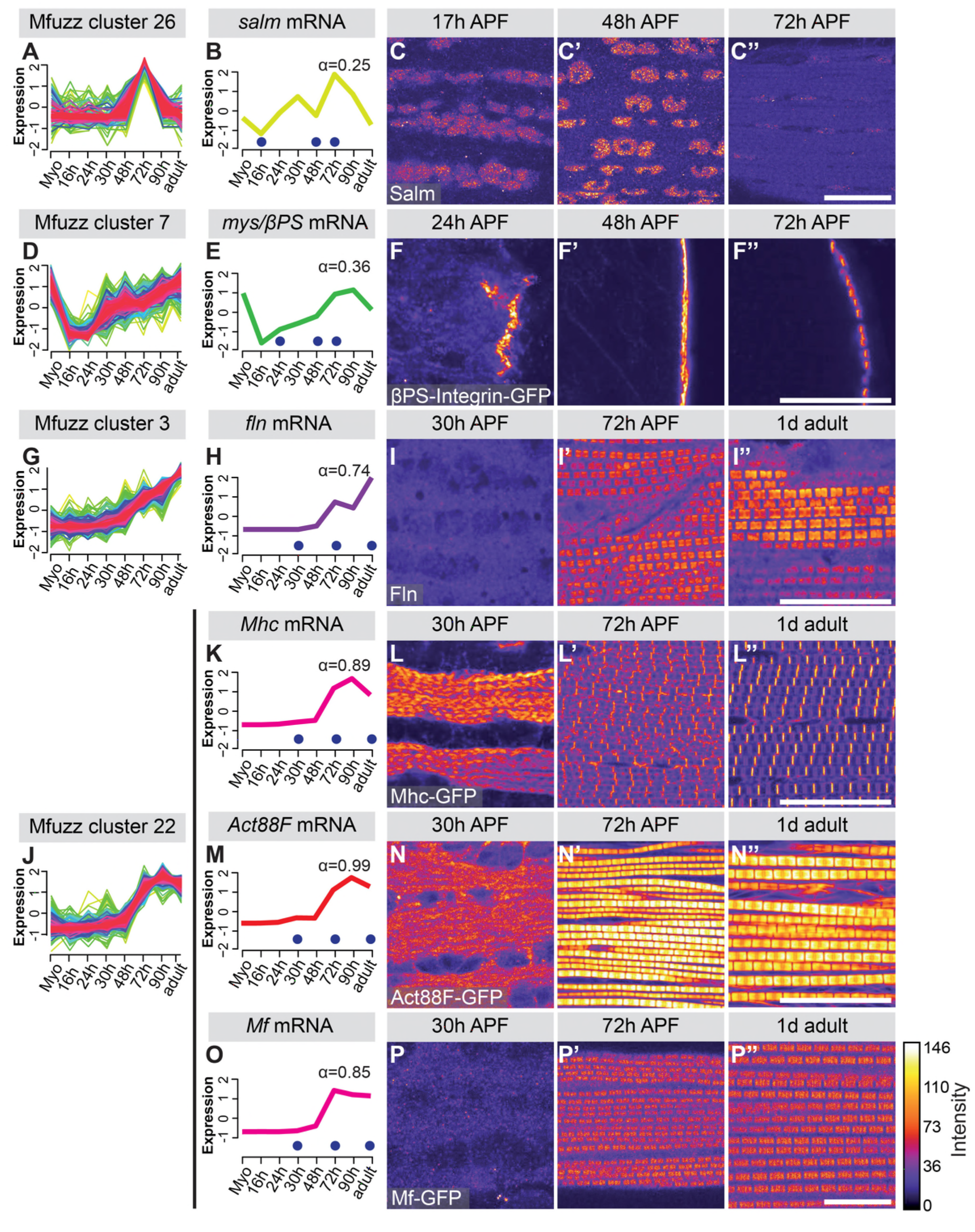
Additional examples of ‘indicator’ gene expression. **(A-P)** Cluster profiles and mRNA dynamics for additional indicator genes. *salm* from cluster 26 (A,B), *mys* (βPS-Integrin) from cluster 7 (D,E), *fln* from cluster 3 (G,H) and *Mhc, Act88F, Mf* from cluster 22 (J,K,M,O) are shown. The respective protein dynamics were visualised with antibodies against the respective protein or GFP fusion protein (C,F,I,L,N,P). Note the high expression of Mhc and Act88F already at 30 h APF, which increases further, while Fln and Mf are only induced to high levels after 30 h APF. Scale bars represent 20 μm.

**Figure 4 - Supplement 1.**
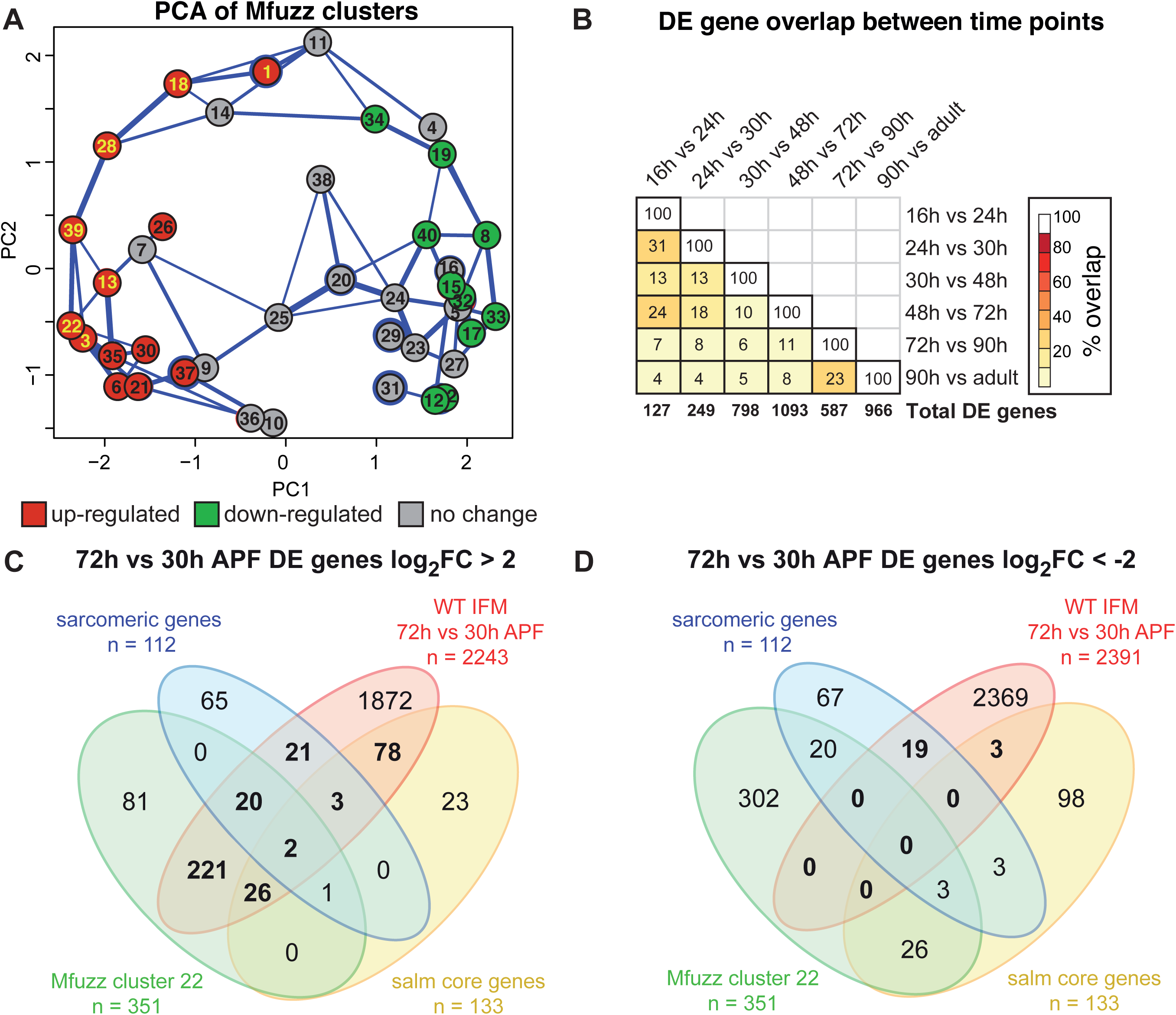
Additional evidence supporting a transition in gene expression between 30 h and 72 h APF. **(A)** Principle component analysis (PCA) of Mfuzz clusters. PC1 largely separates clusters based on 30 h to 72 h dynamics, with significantly “up-regulated” clusters on the left (red) and “down regulated” clusters on the right (green) (see Figure 4F). Sarcomeric and mitochondrial gene clusters shown in yellow script. **(B)** Number of differentially expressed (DE) genes that are the same between time points. Total number of DE genes on bottom. Note that sequential time points share the greatest overlap. **(C,D)** Venn diagrams showing the strong overlap between genes coding for sarcomeric proteins, *salm* core genes, Mfuzz cluster 22 and genes up-regulated (C) but not down-regulated (D) between 30 h and 72 h APF.

**Figure 5 - Supplement 1.**
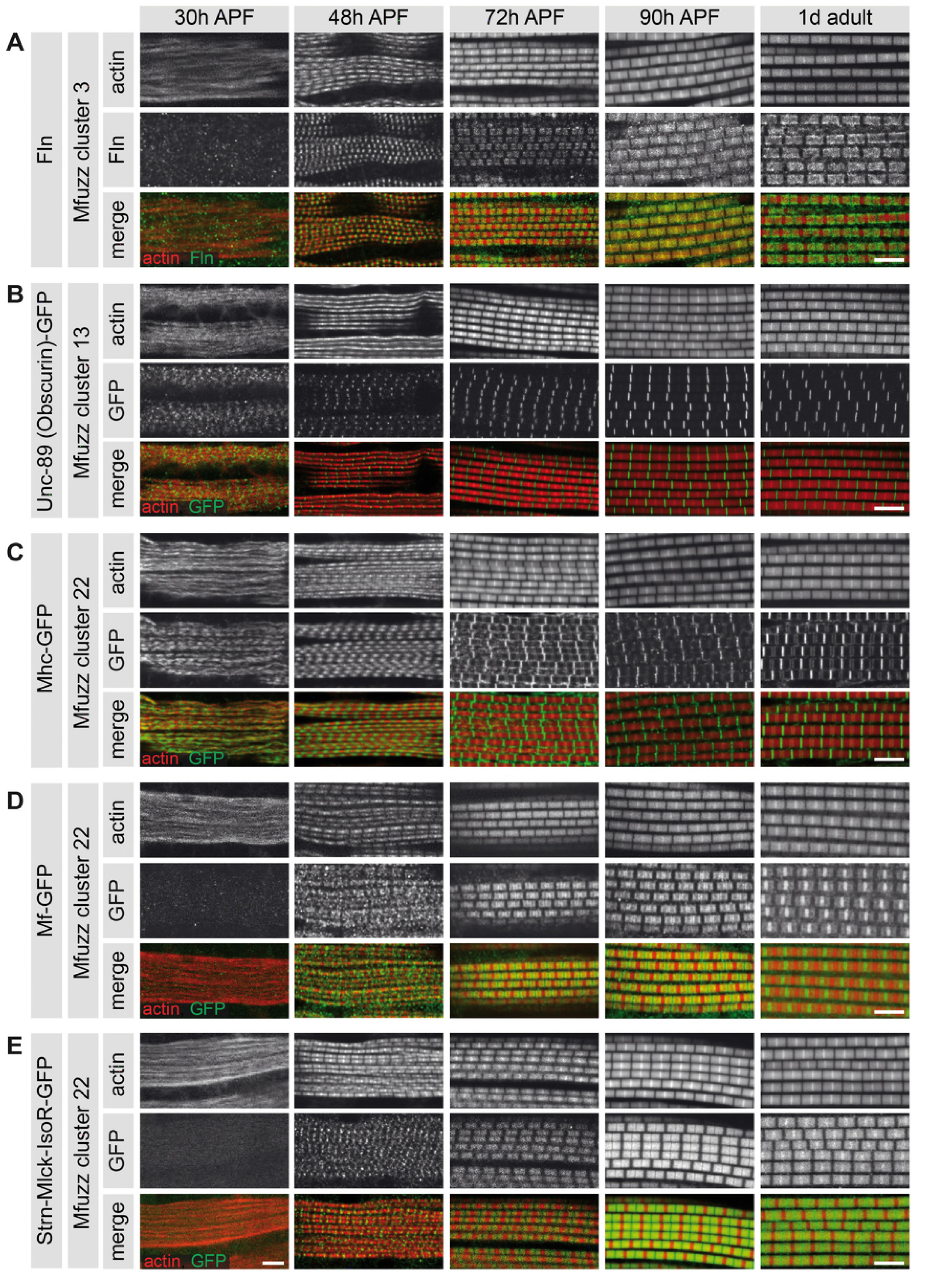
Expression and localisation of thin- and thick-filament structural proteins. **(A-E)** Developing flight muscle myofibrils are stained with phalloidin (red) and the respective sarcomeric proteins (green). Flightin (A) cannot be detected at 30 h, but decorates the thick filament from 48 h APF. Unc-89/Obscurin-GFP (B) labels the M-line from 30 h, and is markedly refined to 72 h APF. Myosin heavy chain (C) is visible in a regular pattern at 30 h APF. Myofilin/Mf (D) cannot be detected at 30 h and decorates the thick filament at 48 h APF. Likewise, IFM-specific Strn-Mlck Isoform R (E) is not expressed at 30 h, but is detected on the thick filament at low levels at 48 h APF. Scale bars represent 5 μm.

**Figure 5 - Movie 1. Twitching time course in developing DLMs**

Confocal movies of spontaneous contraction (twitching) in developing DLMs at 30 h, 36 h, 42 h, 48 h, 60 h and 72 h APF. Fibers are the same as those shown in Figure 5I. Muscle attachments are visualized using a Talin-YPet fusion, which is enriched at the muscle end. Note that fibers weakly contract at 30 h and increase in contraction intensity and frequency until 48 h APF, but then completely stop all contraction by 72 h APF. Scale bar represents 20 μm. Individual movie duration (in seconds) as noted.

**Figure 6 - Supplement 1.**
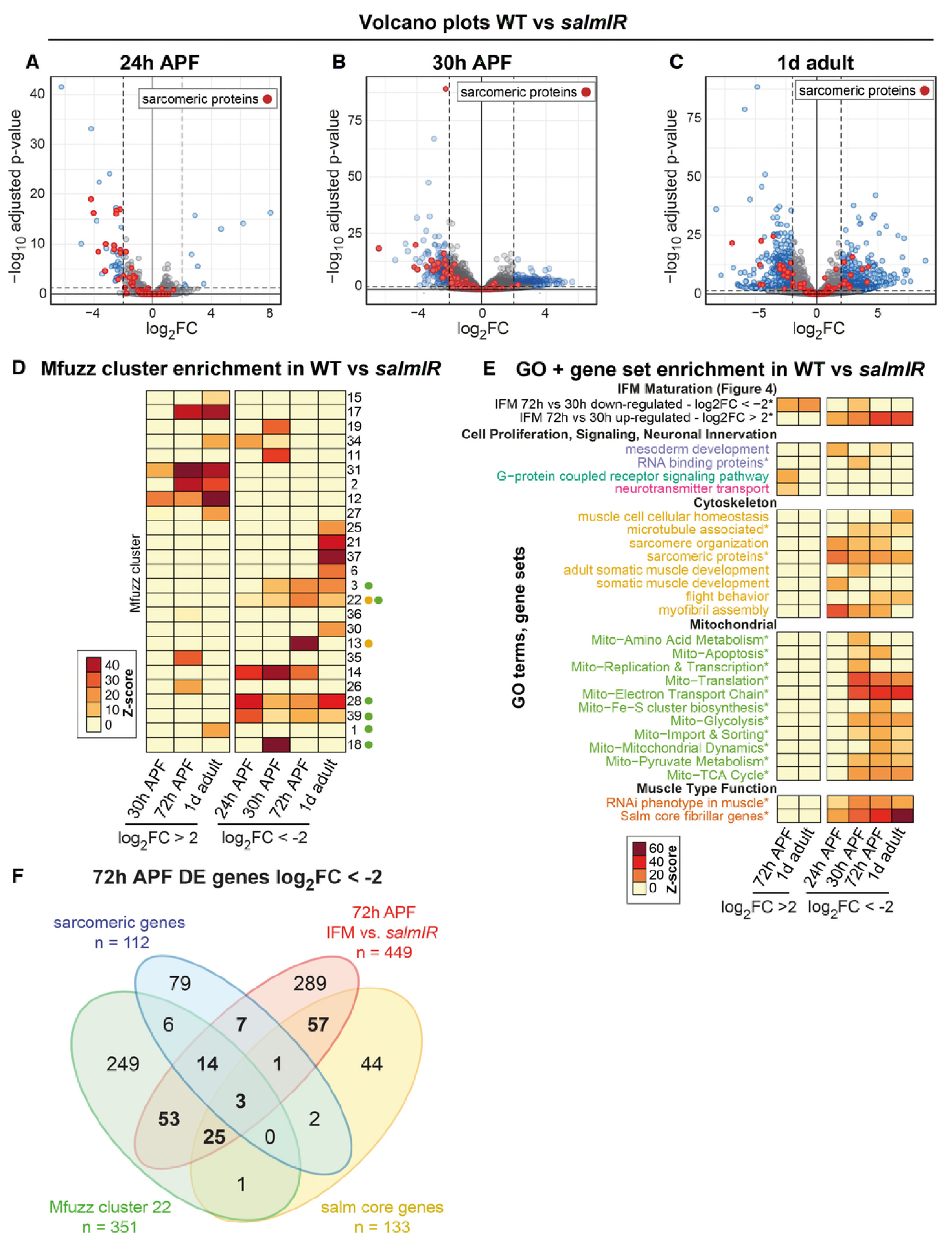
*salm* regulates gene expression during flight muscle development. **(A-C)** Volcano plots of mRNA-Seq data comparing wild-type (WT) versus *salmIR* IFMs at 24h (A), 30h (B) and 1d adult. Significantly differentially expressed (DE) genes (abs(log_2_FC)>2, p < 0.05) are in blue. Note the significant down regulation of genes at all time points, particularly sarcomeric protein coding genes (red). (**D**) Mfuzz cluster enrichment in genes that are DE between WT and *salmIR.* Note that the sarcomeric and mitochondrial clusters (indicated by the green and yellow dots, respectively) are all down-regulated in *salmIR.* Colour scale represents enrichment Z-score. (**E**) GO and gene set enrichment in genes that are DE between WT and *salmIR.* Note that terms down-regulated in *salmIR* are enriched in genes up-regulated in WT from 30 h to 72 h. (**F**) Venn diagram showing the strong overlap between sarcomeric protein coding genes, *salm* core genes, Mfuzz cluster 22 and genes down-regulated in *salmIR* at 72 h APF.

**Figure 6 - Supplement 2.**
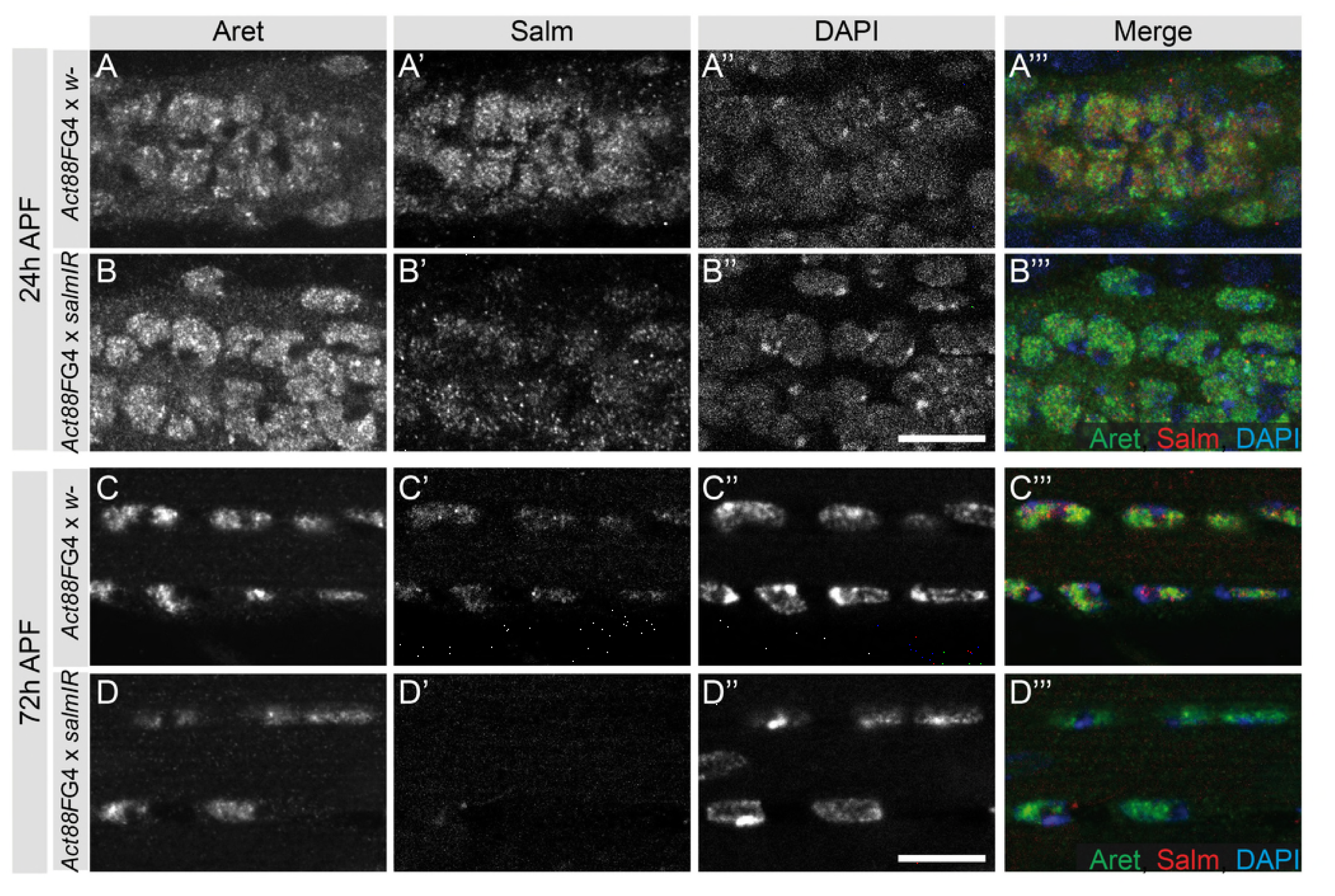
Act88F-GAL4 driven knock-down of *salm* is efficient. **(A-D)** At 24 h APF, both Aret and Salm expression is observed in the nuclei of wild type (A) and *Act88F*≫*salmIR* (B) DLMs. At 72 h APF, Salm protein is present in WT (C) but cannot be detected in the nuclei of *Act88F*≫*salmIR* (D) DLMs. Scale bars represent 10 μm.

**Figure 7 - Movie 2. Twitching in developing *Act88F-GAL4* / + and *Act88F*≫*salmIR* DLMs at 48h APF**

Confocal movies of spontaneous contraction in Act88F-GAL4 / + (control) and *Act88F*≫*salmIR* DLMs at 48h APF. Muscle attachments are visualized using a Talin-YPet fusion. *salmIR* fibers show only a single twitch at 48 h APF, while both single and double twitches are observed in the control. Scale bar represents 20 μm.

**Figure 7 - Movie 3. Twitching in developing Act88F-GAL4 / + and *Act88F*≫*salmIR* DLMs at 72h APF**

72 h APF muscle ends are labelled with a Talin-YPet fusion in Act88F-GAL4 / + (control) and Act88F-GAL4 / *salmIR* DLMs. Control fibers show no contractions at 72 h APF, but *salmIR* fibers continue to contract. Scale bar represents 20 μm.

**Figure 7 - Supplement 1.**
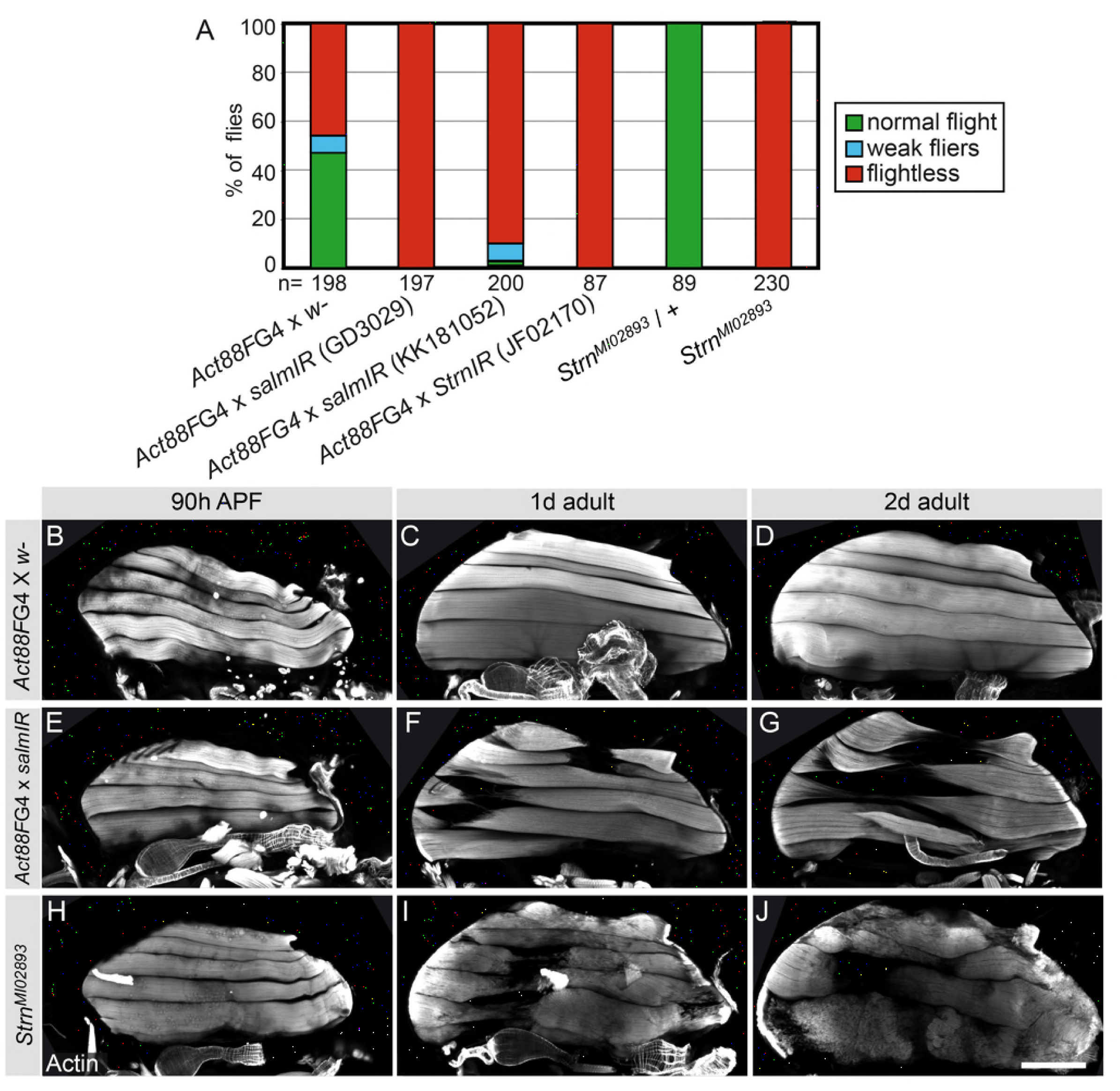
*Act88F*≫*salmIR* and *Strn-Mlck* mutant flies are flightless and IFM fibers rupture in adult flies. **(A)** Adult males from Act88F-GAL4 crossed to two independent *salmIR* hairpins as well as *Strn^M102893^* are flightless. **(B-J)** At 90 h APF, *Act88F-GAL4* / + (B-D),*Act88F*≫*salmIR* (E-G) and *Strn^M102893^* (H-J) all have 6 intact fibers. At 1 day and 2 days after eclosion, both *Act88F*≫*salmIR* (F,G) and *Strn^M102893^* (I,J) show muscle tearing and atrophy. Scale bar represents 200 μm.

**Supplementary Table 1. mRNA-Seq raw data**

The file includes multiple tabs containing the raw or input counts data from bioinformatics analysis, as well as a key to all original data provided in the supplementary tables. This table includes mRNA-Seq counts data, DESeq2 normalized counts data and standard normal counts data used for Mfuzz clustering for wild-type and *salmIR* IFM time points. The averaged core expression profiles for each Mfuzz cluster are also listed.

**Supplementary Table 2. GO-Elite analysis data.**

This table includes multiple tabs containing the GO-Elite analysis of enrichments in Mfuzz clusters as well as genes up- or down-regulated from 30 h - 72 h APF and between wild-type and *salmIR* IFM. It also contains a complete list of all genes included in the ‘User Defined’ gene sets.

**Supplementary Table 3. Summary of sarcomere and myofibril quantifications**

This table includes a numerical summary of quantification values reported graphically in Figures 5, 7 and 8. Quantifications of sarcomere length, myofibril width and myofibril diameter were performed with the MyofibrilJ script (see Materials and Methods). Fiber length and cross-sectional area measurements were performed in Fiji/Image J.

**Supplementary Table 4. DESeq2 pairwise differential expression analysis**

This table contains multiple tabs containing the output data from DESeq2 differential expression analysis between sequential IFM development time points, from 30-72h APF as well as between WT and *salmIR* IFM.

## References

Anant, S., Roy, S., and VijayRaghavan, K. (1998). Twist and Notch negatively regulate adult muscle differentiation in Drosophila. Development 125, 1361.

Asano, T., Ishizuka, T., Morishima, K., and Yawo, H. (2015). Optogenetic induction of contractile ability in immature C2C12 myotubes. Sci Rep 5, 8317.

Bate, M., Rushton, E., and Currie, D.A. (1991). Cells with persistent twist expression are the embryonic precursors of adult muscles in Drosophila. Development 113, 79–89.

Brinegar, A.E., Xia, Z., Loehr, J.A., Li, W., Rodney, G.G., and Cooper, T.A. (2017). Extensive alternative splicing transitions during postnatal skeletal muscle development are required for calcium handling functions. eLife 6, 399.

Bryantsev, A.L., Baker, P.W., Lovato, T.L., Jaramillo, M.S., and Cripps, R.M. (2012). Differential requirements for Myocyte Enhancer Factor-2 during adult myogenesis in Drosophila. Developmental Biology 361, 191–207.

Bullard, B., and Pastore, A. (2011). Regulating the contraction of insect flight muscle. J Muscle Res Cell Motil 32, 303–313.

Chen, H. (2016). VennDiagram: Generate High-Resolution Venn and Euler Plot.

Clark, I.E., Dodson, M.W., Jiang, C., Cao, J.H., Huh, J.R., Seol, J.H., Yoo, S.J., Hay, B.A., and Guo, M. (2006). Drosophila pink1 is required for mitochondrial function and interacts genetically with parkin. Nature 441, 1162–1166.

Dobin, A., Davis, C.A., Schlesinger, F., Drenkow, J., Zaleski, C., Jha, S., Batut, P., Chaisson, M., and Gingeras, T.R. (2013). STAR: ultrafast universal RNA-seq aligner. Bioinformatics 29, 15–21.

Du, A., Sanger, J.M., and Sanger, J.W. (2008). Cardiac myofibrillogenesis inside intact embryonic hearts. Developmental Biology 318, 236–246.

Dutta, D., Anant, S., Ruiz-Gómez, M., Bate, M., and VijayRaghavan, K. (2004). Founder myoblasts and fibre number during adult myogenesis in Drosophila. Development 131, 3761–3772.

Ehler, E., and Gautel, M. (2008). The sarcomere and sarcomerogenesis. Adv. Exp. Med. Biol. 642, 1–14.

Fernandes, J., Bate, M., and VijayRaghavan, K. (1991). Development of the indirect flight muscles of Drosophila. Development 113, 67–77.

Filardo, P., and Ephrussi, A. (2003). Bruno regulates gurken during Drosophila oogenesis. Mechanisms of Development 120, 289–297.

Förster, D., and Luschnig, S. (2012). Src42A-dependent polarized cell shape changes mediate epithelial tube elongation in Drosophila. Nature Cell Biology 14, 526–534.

Futschik, M.E., and Carlisle, B. (2005). Noise-robust soft clustering of gene expression time-course data. J Bioinform Comput Biol 3, 965–988.

Gajewski, K.M., and Schulz, R.A. (2010). CF2 represses Actin 88F gene expression and maintains filament balance during indirect flight muscle development in Drosophila. PLoS ONE 5, e10713.

García-Zaragoza, E., Mas, J.A., Vivar, J., Arredondo, J.J., and Cervera, M. (2008). CF2 activity and enhancer integration are required for proper muscle gene expression in Drosophila. Mechanisms of Development 125, 617–630.

Gautel, M., and Djinovic-Carugo, K. (2016). The sarcomeric cytoskeleton: from molecules to motion. J Exp Biol 219, 135–145.

Gokhin, D.S., and Fowler, V.M. (2013). A two-segment model for thin filament architecture in skeletal muscle. Nature Reviews Molecular Cell Biology 14, 113–119.

Goodman, J.W. (1968). Introduction to Fourier Optics Fourier Optics.

Gu, Z., Eils, R., and Schlesner, M. (2016). Complex heatmaps reveal patterns and correlations in multidimensional genomic data. Bioinformatics 32, 2847–2849.

Hinits, Y., and Hughes, S.M. (2007). Mef2s are required for thick filament formation in nascent muscle fibres. Development 134, 2511–2519.

Josephson, R. (2006). Comparative Physiology of Insect Flight Muscle. In Nature’s Versatile Engine: Insect Flight Muscle Inside and Out, J. Vigoreaux, ed. (Georgetown, TX: Landes Bioscience)), pp. 35–43.

Kelly, K.K., Meadows, S.M., and Cripps, R.M. (2002). Drosophila MEF2 is a direct regulator of Actin57B transcription in cardiac, skeletal, and visceral muscle lineages. Mechanisms of Development 110, 39–50.

Kumar, L., and E Futschik, M. (2007). Mfuzz: a software package for soft clustering of microarray data. Bioinformation 2, 5–7.

Kühnlein, R.P., Frommer, G., Friedrich, M., Gonzalez-Gaitan, M., Weber, A., WagnerBernholz, J.F., Gehring, W.J., Jäckle, H., and Schuh, R. (1994). spalt encodes an evolutionarily conserved zinc finger protein of novel structure which provides homeotic gene function in the head and tail region of the Drosophila embryo. The EMBO Journal 13, 168–179.

Lange, S., Ehler, E., and Gautel, M. (2006). From A to Z and back? Multicompartment proteins in the sarcomere. Trends in Cell Biology 16, 11–18.

Leiss, D., Hinz, U., Gasch, A., Mertz, R., and Renkawitz-Pohl, R. (1988). Beta 3 tubulin expression characterizes the differentiating mesodermal germ layer during Drosophila embryogenesis. Development 104, 525–531.

Lemke, S.B., and Schnorrer, F. (2017a). Mechanical forces during muscle development. Mechanisms of Development 144, 92–101.

Lemke, S.B., and Schnorrer, F. (2017b). In Vivo Imaging of Muscle-tendon Morphogenesis in Drosophila Pupae. JoVE.

Li, H., Handsaker, B., Wysoker, A., Fennell, T., Ruan, J., Homer, N., Marth, G., Abecasis, G., Durbin, R., 1000 Genome Project Data Processing Subgroup (2009). The Sequence Alignment/Map format and SAMtools. Bioinformatics 25, 2078–2079.

Liao, Y., Smyth, G.K., and Shi, W. (2014). featureCounts: an efficient general purpose program for assigning sequence reads to genomic features. Bioinformatics 30, 923–930.

Liotta, D., Han, J., Elgar, S., Garvey, C., Han, Z., and Taylor, M.V. (2007). The Him Gene Reveals a Balance of Inputs Controlling Muscle Differentiation in Drosophila. Current Biology 17, 1409–1413.

Llewellyn, M., Barretto, R., Delp, S., and Schnitzer, M. (2008). Minimally invasive highspeed imaging of sarcomere contractile dynamics in mice and humans. Nature 454, 784788.

Love, M.I., Huber, W., and Anders, S. (2014). Moderated estimation of fold change and dispersion for RNA-seq data with DESeq2. Genome Biology 15, 550.

Marty, F., Rockel-Bauer, C., Simigdala, N., Brunner, E., and Basler, K. (2014). Large-scale imaginal disc sorting: A protocol for “omics-”approaches. Methods 68, 260–264.

Mazelet, L., Parker, M.O., Li, M., Arner, A., and Ashworth, R. (2016). Role of Active Contraction and Tropomodulins in Regulating Actin Filament Length and Sarcomere Structure in Developing Zebrafish Skeletal Muscle. Front Physiol 7, 91.

Montfort, J., Le Cam, A., Gabillard, J.-C., and Rescan, P.-Y. (2016). Gene expression profiling of trout regenerating muscle reveals common transcriptional signatures with hyperplastic growth zones of the post-embryonic myotome. BMC Genomics 17, 810.

Neuwirth, E. (2015). Package “RColorBrewer.” 1–5.

Orfanos, Z., and Sparrow, J.C. (2013). Myosin isoform switching during assembly of the Drosophila flight muscle thick filament lattice. Journal of Cell Science 126, 139–148.

Orfanos, Z., Leonard, K., Elliott, C., Katzemich, A., Bullard, B., and Sparrow, J. (2015). Sallimus and the Dynamics of Sarcomere Assembly in Drosophila Flight Muscles. Journal of Molecular Biology 427, 2151–2158.

Pérez-Moreno, J.J., Bischoff, M., Martín-Bermudo, M.D., and Estrada, B. (2014). The conserved transmembrane proteoglycan Perdido/Kon-tiki is essential for myofibrillogenesis and sarcomeric structure in Drosophila. Journal of Cell Science 127, 3162–3173.

Potthoff, M.J., Arnold, M.A., McAnally, J., Richardson, J.A., Bassel-Duby, R., and Olson, E.N. (2007). Regulation of skeletal muscle sarcomere integrity and postnatal muscle function by Mef2c. Molecular and Cellular Biology 27, 8143–8151.

Racca, A.W., Klaiman, J.M., Pioner, J.M., Cheng, Y., Beck, A.E., Moussavi-Harami, F., Bamshad, M.J., and Regnier, M. (2016). Contractile properties of developing human fetal cardiac muscle. J. Physiol. (Lond.) 594, 437–452.

Reedy, M.C., and Beall, C. (1993). Ultrastructure of developing flight muscle in Drosophila. I. Assembly of myofibrils. Developmental Biology 160, 443–465.

Reedy, M., Bullard, B., and Vigoreaux, J. (2000). Flightin is essential for thick filament assembly and sarcomere stability in Drosophila flight muscles. Journal of Cell Biology 151, 1483.

Regev, G.J., Kim, C.W., Tomiya, A., Lee, Y.P., Ghofrani, H., Garfin, S.R., Lieber, R.L., and Ward, S.R. (2011). Psoas Muscle Architectural Design, In Vivo Sarcomere Length Range, and Passive Tensile Properties Support Its Role as a Lumbar Spine Stabilizer. Spine 36, E1666–E1674.

Roper, K., Mao, Y., and Brown, N. (2005). Contribution of sequence variation in Drosophila actins to their incorporation into actin-based structures in vivo. Journal of Cell Science 118, 3937.

Sandmann, T., Jensen, L., Jakobsen, J., Karzynski, M., Eichenlaub, M., Bork, P., and Furlong, E. (2006). A temporal map of transcription factor activity: mef2 directly regulates target genes at all stages of muscle development. Developmental Cell 10, 797807.

Sanger, J.W., Wang, J., Fan, Y., White, J., Mi-Mi, L., Dube, D.K., Sanger, J.M., and Pruyne, D. (2017). Assembly and Maintenance of Myofibrils in Striated Muscle. Handb Exp Pharmacol 235, 39–75.

Sanger, J.W., Wang, J., Holloway, B., Du, A., and Sanger, J.M. (2009). Myofibrillogenesis in skeletal muscle cells in zebrafish. Cell Motil. Cytoskeleton 66, 556–566.

Sarov, M., Barz, C., Jambor, H., Hein, M.Y., Schmied, C., Suchold, D., Stender, B., Janosch, S., KJ, V.V., Krishnan, R.T., et al. (2016). A genome-wide resource for the analysis of protein localisation in Drosophila. eLife 5, 2185.

Schindelin, J., Arganda-Carreras, I., Frise, E., Kaynig, V., Longair, M., Pietzsch, T., Preibisch, S., Rueden, C., Saalfeld, S., Schmid, B., et al. (2012). Fiji: an open-source platform for biological-image analysis. Nature Methods 9, 676–682.

Schnorrer, F., Kalchhauser, I., and Dickson, B. (2007). The transmembrane protein Kon-tiki couples to Dgrip to mediate myotube targeting in Drosophila. Developmental Cell 12, 751–766.

Schnorrer, F., Schönbauer, C., Langer, C.C.H., Dietzl, G., Novatchkova, M., Schernhuber, K., Fellner, M., Azaryan, A., Radolf, M., Stark, A., et al. (2010). Systematic genetic analysis of muscle morphogenesis and function in Drosophila. Nature 464, 287–291.

Schönbauer, C., Distler, J., Jährling, N., Radolf, M., Dodt, H.-U., Frasch, M., and Schnorrer, F. (2011). Spalt mediates an evolutionarily conserved switch to fibrillar muscle fate in insects. Nature 479, 406–409.

Sevdali, M., Kumar, V., Peckham, M., and Sparrow, J. (2013). Human congenital myopathy actin mutants cause myopathy and alter Z-disc structure in Drosophila flight muscle. Neuromuscular Disorders 23, 243–255.

Shwartz, A., Dhanyasi, N., Schejter, E.D., and Shilo, B.-Z. (2016). The Drosophila formin Fhos is a primary mediator of sarcomeric thin-filament array assembly. eLife 5, D786.

Soler, C., and Taylor, M. (2009). The Him gene inhibits the development of Drosophila flight muscles during metamorphosis. Mechanisms of Development 126, 595–603.

Soler, C., Han, J., and Taylor, M.V. (2012). The conserved transcription factor Mef2 has multiple roles in adult Drosophila musculature formation. Development 139, 1270–1275.

Sparrow, J., and Schöck, F. (2009). The initial steps of myofibril assembly: integrins pave the way. Nature Reviews Molecular Cell Biology 10, 293–298.

Spletter, M.L., and Schnorrer, F. (2014). Transcriptional regulation and alternative splicing cooperate in muscle fiber-type specification in flies and mammals. Experimental Cell Research 321, 90–98.

Spletter, M.L., Barz, C., Yeroslaviz, A., Schönbauer, C., Ferreira, I.R.S., Sarov, M., Gerlach, D., Stark, A., Habermann, B.H., and Schnorrer, F. (2015). The RNA-binding protein Arrest (Bruno) regulates alternative splicing to enable myofibril maturation in Drosophila flight muscle. EMBO Rep 16, 178–191.

Stronach, B.E., Renfranz, P.J., Lilly, B., and Beckerle, M.C. (1999). Muscle LIM proteins are associated with muscle sarcomeres and require dMEF2 for their expression during Drosophila myogenesis. Molecular Biology of the Cell 10, 2329–2342.

Syme, D., and Josephson, R. (2002). How to build fast muscles: synchronous and asynchronous designs. Integrative and Comparative Biology 42, 762.

Tanaka, K.K.K., Bryantsev, A.L., and Cripps, R.M. (2008). Myocyte enhancer factor 2 and chorion factor 2 collaborate in activation of the myogenic program in Drosophila. Molecular and Cellular Biology 28, 1616–1629.

Tskhovrebova, L., and Trinick, J. (2003). Titin: properties and family relationships. Nature Reviews Molecular Cell Biology 4, 679–689.

Vigoreaux, J.O. (2006). Molecular Basis of Muscle Structure. In Muscle Development in Drosophila, (New York, NY: Springer New York), pp. 143–156.

Wallgren-Pettersson, C., Sewry, C.A., Nowak, K.J., and Laing, N.G. (2011). Nemaline myopathies. Semin Pediatr Neurol 18, 230–238.

Warren, G.L., Summan, M., Gao, X., Chapman, R., Hulderman, T., and Simeonova, P.P. (2007). Mechanisms of skeletal muscle injury and repair revealed by gene expression studies in mouse models. J. Physiol. (Lond.) 582, 825–841.

Weitkunat, M., and Schnorrer, F. (2014). A guide to study Drosophila muscle biology. Methods 68, 2–14.

Weitkunat, M., Brasse, M., Bausch, A.R., and Schnorrer, F. (2017). Mechanical tension and spontaneous muscle twitching precede the formation of cross-striated muscle in vivo. Development 144, 1261–1272.

Weitkunat, M., Kaya-Copur, A., Grill, S.W., and Schnorrer, F. (2014). Tension and force-resistant attachment are essential for myofibrillogenesis in Drosophila flight muscle. Curr Biol 24, 705–716.

Wickham, H. (2009). ggplot2: Elegant Graphics for Data Analysis (Springer).

Wickham, H. (2007). Reshaping Data with the reshapePackage. Journal of Statistical Software 21, 1–20.

Wickham, H. (2011). The Split-Apply-Combine Strategy for Data Analysis. Journal of Statistical Software 40, 1–29.

Zambon, A.C., Gaj, S., Ho, I., Hanspers, K., Vranizan, K., Evelo, C.T., Conklin, B.R., Pico, A.R., and Salomonis, N. (2012). GO-Elite: a flexible solution for pathway and ontology over-representation. Bioinformatics 28, 2209–2210.

Zappia, M.P., and Frolov, M.V. (2016). E2F function in muscle growth is necessary and sufficient for viability in Drosophila. Nature Communications 7, 10509.

Zhao, Y., Li, J., Liu, H., Xi, Y., Xue, M., Liu, W., Zhuang, Z., and Lei, M. (2015). Dynamic transcriptome profiles of skeletal muscle tissue across 11 developmental stages for both Tongcheng and Yorkshire pigs. BMC Genomics 16, 377.

Zheng, Q., Zhang, Y., Chen, Y., Yang, N., Wang, X.-J., and Zhu, D. (2009). Systematic identification of genes involved in divergent skeletal muscle growth rates of broiler and layer chickens. BMC Genomics 10, 87.

